# Yeast Rgd3 is a phospho-regulated F-BAR-containing RhoGAP involved in the regulation of Rho3 distribution and cell morphology

**DOI:** 10.1101/2020.05.04.077206

**Authors:** Robert M. Gingras, Kyaw Myo Lwin, Abigail M. Miller, Anthony Bretscher

## Abstract

Polarized growth requires the integration of polarity pathways with the delivery of exocytic vesicles for cell expansion and counterbalancing endocytic uptake. In budding yeast, the myosin-V Myo2 is aided by the kinesin-related protein Smy1 in carrying out the essential Sec4-dependent transport of secretory vesicles to sites of polarized growth. Over-expression suppressors of a conditional *myo2 smy1* mutant identified a novel F-BAR-containing RhoGAP, Rgd3, that has activity primarily on Rho3, but also Cdc42. Internally tagged Rho3 is restricted to the plasma membrane in a gradient corresponding to cell polarity that is altered upon Rgd3 over-expression. Rgd3 itself is localized to dynamic polarized vesicles that, while distinct from constitutive secretory vesicles, are dependent on actin and Myo2 function. *In vitro* Rgd3 associates with liposomes in a PIP_2_-enhanced manner. Further, the Rgd3 C-terminal region contains several phosphorylatable residues within a reported SH3-binding motif. An unphosphorylated mimetic construct is active and highly polarized, while the phospho-mimetic form is not. Rgd3 is capable of activating Myo2, dependent on its phospho-state and Rgd3 overexpression rescues aberrant Rho3 localization and cell morphologies seen at the restrictive temperature in the *myo2 smy1* mutant. We propose a model where Rgd3 functions to modulate and maintain Rho3 polarity during growth.

## Introduction

Budding yeast grows in a polarized manner with the cell selecting one site for bud emergence, followed by bud growth, organelle segregation, and ultimately cytokinesis. Achieving polarized growth requires the integration of signaling pathways and the assembly of a polarized cytoskeleton that guides membrane and organelle transport. In budding yeast, the microtubule cytoskeleton is responsible for nuclear orientation and mitosis, and the actin-based cytoskeleton is used for the transport and segregation of organelles, endocytosis, and the contractile ring during cytokinesis (reviewed in Schott et al., 2002).

The transport of post-Golgi secretory vesicles and segregation of all organelles is dependent on the assembly of actin cables at the bud cortex and bud neck by the two formins, Bni1 and Bnr1, respectively. These cables serve as tracks for the two myosin-V motors in yeast, the essential Myo2 that transports secretory vesicles, mitochondria, peroxisomes, the vacuole and the cytoplasmic microtubules for nuclear orientation, and Myo4 that transports mRNAs and the endoplasmic reticulum. Myo2 is a typical myosin-V heavy chain consisting of an N-terminal motor domain followed by IQ-motif repeats, a coiled-coil dimerization domain, and terminating in a large globular cargo-binding tail domain. The tail domain associates with organelle-specific receptors to mediate their transport. Here it associates with the Rab protein Sec4 for transport of secretory vesicles, Mmr1 and Ypt11 for mitochondrial transport, Inp2 for peroxisome transport, Vac17 for vacuole transport, and Kar9 for association with microtubule plus ends (reviewed in (Hammer and Sellers, 2012).

The transport of secretory vesicles by Myo2 is an essential process which has been shown genetically to involve the non-essential kinesin-related protein Smy1 (Lillie and Brown, 1994, 1992). Smy1 interacts directly with Myo2, and we have previously shown that it functions to specifically enhance the association of Myo2 with the secretory vesicle receptor, Sec4 (Lwin et al., 2016; Lillie and Brown, 1994). In this report, we employ a genetic approach to uncover additional components that may be involved in the direct or indirect regulation of secretory vesicle transport. This screen led to the discovery of a previously uncharacterized F-BAR-containing RhoGAP (GTPase Activating Protein), which we named Rgd3, that regulates Rho3 and Cdc42. Interestingly, Rgd3 associates with vesicles polarized to sites of growth which are distinct from constitutive secretory vesicles. Furthermore, the polarity of Rgd3 vesicles is dependent on an intact actin cytoskeleton. Our study shows that Rgd3 is capable of redistributing Rho3 on the plasma membrane and contributes to overall cell morphology, particularly in cells where polarity is compromised.

## Results

### Identification of proteins whose overexpression suppresses conditional *myo2 smy1* mutants

An essential function of the yeast myosin-V motor encoded by *MYO2* is to transport secretory vesicles to the site of cell growth (Johnston et al., 1991; Schott et al., 1999). The kinesin-related protein Smy1 functions with Myo2 to enhance the association of Myo2 with its receptor on secretory vesicles, the Rab Sec4 (Beningo et al., 2000; Lillie and Brown, 1994; Lwin et al., 2016). We have described *myo2* mutant alleles partially compromised in binding Sec4 in which *SMY1* becomes essential and used these alleles to generate *myo2 smy1* conditional mutations (Figure 1A and B). In order to explore what ancillary factors might function directly or indirectly in this process, we used two different *myo2 smy1* strains to screen our regulated *GAL1*-cDNA overexpression library (Liu et al., 1992) for clones that could suppress the temperature sensitivity at 35°. After isolating and retesting the plasmids from about 250,000 total transformants, 25 clones were recovered and sequenced to yield cDNAs from 9 different genes (Figure 1C). Among the cDNAs recovered were: *SMY1* itself; genes encoding the Rab proteins Sec4 and Ypt31/32, both of which interact with the cargo-binding domain of Myo2 (Lipatova et al., 2008; Santiago-Tirado et al., 2011); and genes encoding Cmd1 and Mlc1, both of which associate with the lever arm of Myo2 (Brockerhoff et al., 1994; Stevens and Davis, 1998). In addition, we recovered an uncharacterized gene, *YHR182W,* that has a putative RhoGAP domain, and *SMI1* (aka *KNR4*), that has been proposed to play a role in the regulation of cell wall synthesis (Basmaji et al., 2005; Roumanie et al., 2001). When genomic DNA was recovered to express the genes from their own promoters on high-copy 2μ plasmids, the identified genes all suppressed the conditional growth defect of *myo2-41 smy1-15* at 35°C (Figure 1D). However, only four—*YHR182W, MLC1, SMY1,* and *YPT31*—suppressed *myo2-57 smy1-12*, presumably due to the more severe mutation within the Rab binding site in the Myo2-cargo-binding domain (Figure 1B and S1A).

**Figure 1.**
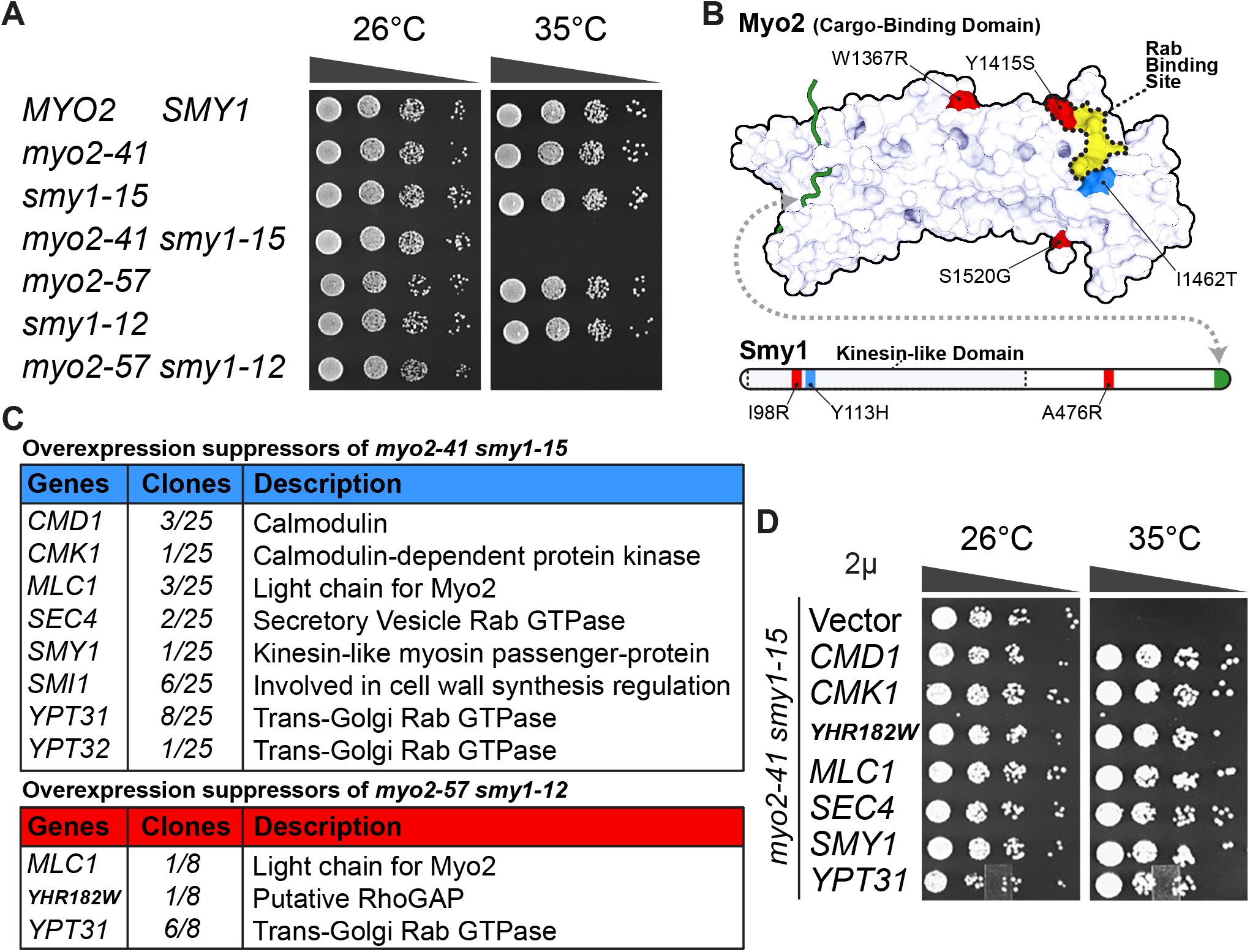
Screen for overexpression suppressors of *myo2 smy1* conditional mutants. **A.** Growth of the temperature-sensitive strains *myo2-41 smy1-15* and *myo2-57 smy1-12.* Ten-fold serial dilutions grown on YPD at either 26°C or 35°C for two days. **B.** Mutations in *myo2-41 smy1-15* (blue) and *myo2-57 smy1-12* (red) compared to the Myo2-Rab binding site (yellow/dashed outline) and the Myo2-Smy1 binding site (green). PDB: 6IXQ, Tang et al., 2019. **C.** Summary table of Gal-overexpression suppressors identified in both strains. **D.** Confirmation of overexpression suppression of the identified cDNAs on 2μ vectors in *myo2-41 smy1-15*.

### *YHR182W*, now *RGD3*, encodes a RhoGAP for Rho3 and Cdc42

Previous studies have identified nine RhoGAP proteins in yeast, Bem2, Bem3, Bag7, Lrg1, Sac7, Rgd1, Rgd2, Rga1, and Rga2. *YHR182W* represents one of two additional genes (the other being *ECM25*) whose protein products were computationally predicted to have RhoGAP activity (Roumanie et al., 2001). By sequence analysis, *YHR182W* is most closely related to *RGD1* and *RGD2* (about 15% identity, 25% similarity), thus, for consistency, we decided to name this third relative *RGD3*. All three protein sequences were subjected to HHpred homology detection and found to have a putative N-terminal F-BAR (Fes/CIP4 homology-Bin-Amphiphysin-Rvs) domain and a C-terminal GAP domain (Zimmermann et al., 2018). Rgd2 and Rgd3 also have a DEP (Dishevelled, Egl-10, Pleckstrin-homology) domain—generally involved in membrane-targeting and G-protein signaling (Burchett, 2002; Wong et al., 2000)—embedded in the F-BAR domain (Figure 2A). The putative F-BAR and DEP domains of Rgd3 were not previously identified or annotated within the Pfam or SMART databases. These three are the only RhoGAP proteins in *S. cerevisiae* with F-BAR domains and together appear to be distantly structurally related to human GMIP and srGAPs.

**Figure 2.**
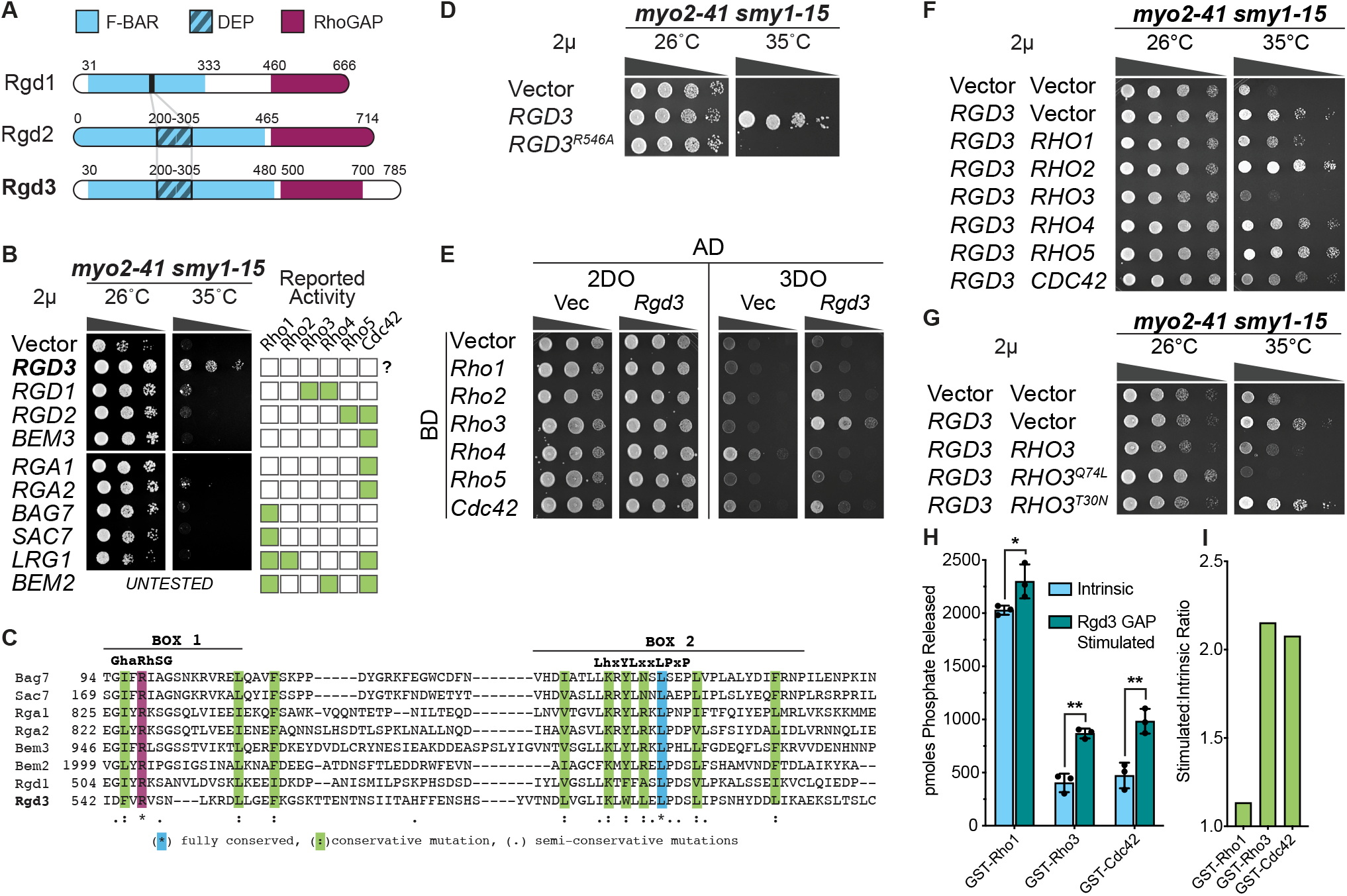
Rgd3 (*YHR182W*) is a RhoGAP for Rho3 and Cdc42. **A.** Schematic diagram of domains in Rgd3 and related RhoGAPs in *S. cerevisiae* as determined by HHPred. **B.** Rgd3 is the only RhoGAP that suppresses *myo2-41 smy1-15* temperature sensitivity. Activities reported in the literature are shown alongside. **C.** Alignment of the Rgd3 RhoGAP active site with other known RhoGAPs in *S. cerevisiae*. **D.** An arginine residue, that aligns with other known catalytic arginine-finger motifs, is required for Rgd3’s suppression of *myo2-41 smy1-15* temperature sensitivity. **E.** Rgd3 interacts strongly with Rho3 and weakly with Cdc42 via yeast two-hybrid. 2DO = two amino acid (Leu, Trp) drop-out; 3DO = three amino acid (Leu, Trp, His) drop-out. **F.** Rho3 2μ overexpression exhibits the greatest counter-suppression of temperature sensitivity in *myo2-41 smy1-15 2μ RGD3.* Rho proteins were introduced on a second, separate 2μ plasmid. **G.** Constitutively active Rho3 (*RHO3*^*Q74L*^), and not the inactive form (*RHO3^T30N^*), exhibits the same counter-suppression of Rgd3 suppression seen in F. **H.** Rgd3’s GAP-domain shows greater GAP activity against Rho3 and Cdc42 than Rho1 in a colorimetric malachite green GTPase assay, *n*=3. *, p≤0.05; **, p≤0.005 by Student’s T-test. **I.** Ratio of intrinsic GTP hydrolysis to Rgd3-stimulated hydrolysis from H.

To test if the ability to suppress *myo2-41 smy1-12* is shared by other yeast RhoGAP proteins, we overexpressed eight of the RhoGAPs from their own promoters on 2μ plasmids. Of all the RhoGAPs, only *RGD3* overexpression could confer suppression, indicating that it plays a specific role distinct from all the other RhoGAP proteins tested (Figure 2B).

RhoGAP domains have a critical arginine residue that facilitates the hydrolysis of GTP to GDP and Pi (Bourne, 1997). Clustal alignment of Rgd3 with seven yeast RhoGAP proteins identified residue R546 as the critical arginine in Rgd3 (Figure 2C) (Madeira et al., 2019). Overexpression of *RGD3*^*R546A*^ failed to suppress the *myo2-14 smy1-15* conditional mutant, showing that GAP activity is necessary for its function in suppression (Figure 2D and S2A).

Both genetic and biochemical approaches were used to identify which of the six yeast Rho proteins (Cdc42, Rho1-5) is a substrate for Rgd3. Two-hybrid analysis revealed the strongest interaction with Rho3, and a weaker interaction with Cdc42 (Figure 2E). As RhoGAPs are negative regulators of Rho proteins, we also tested if overexpression of any of the Rho proteins could abolish the ability of *RGD3* overexpression to suppress the conditional *myo2-14 smy1-15* mutant. Once again, Rho3 was identified as having the greatest effect, but an effect was also seen with Cdc42 (Figure 2F). The same result was observed in *myo2-57 smy1-12* (Figure S1B). This effect of *RHO3* overexpression was dependent on its GTPase activity, as a constitutively active *RHO3*^*Q74L*^ had a stronger effect than wild-type *RHO3* or the dominant negative *RHO3*^*T30N*^, which showed no effect (Figure 2G). These genetic data suggest that Rho3, and possibly also Cdc42, are substrates for Rgd3. Of the six Rho proteins, only Cdc42, Rho1, and Rho3 are either essential, or nearly essential. We therefore assayed the ability of Rgd3 to stimulate the hydrolysis of GTP by these three Rho proteins loaded with GTP. Under the conditions used, Rgd3 was able to stimulate the GTPase activities of Rho3 and Cdc42 about two-fold but had minimal effect on Rho1 (Figure 2H and I). Our genetic and biochemical data therefore indicate that Rgd3 is a GAP for Rho3, and likely Cdc42.

Deletion or overexpression of *RGD3* does not confer any growth phenotype in otherwise wild-type yeast (Figure S1C). Moreover, it shows no synthetic lethality with *rgd1*Δ, the only other RhoGAP that has been reported to function with Rho3 (Figure S3A and B). Surprisingly, *rgd3*Δ cells, *rgd1*Δ cells, and *rgd3Δ rgd1*Δ cells all appear to have an unimpaired actin cytoskeleton (Figure S3C).

### Rgd3 modulates the distribution of Rho3 on the plasma membrane

As Rgd3 is a RhoGAP for Rho3, we wished to localize Rho3 in living cells. It has been notoriously difficult to define the *in vivo* location of fully functional Rho proteins by tagging as the small GTPases are C-terminally prenylated and occasionally N-terminally palmitoylated (Ren et al., 2008; Wu and Brennwald, 2010). Rho3 is an example which is modified at both termini, precluding traditional methods of fluorescent-tagging. For this reason, Rho3 has been localized by immunofluorescence (Wu and Brennwald, 2010), but no functionally tagged version has been described. We therefore assessed the effect of inserting fluorescent proteins internally at several locations. In just one case, Rho3 was functional when mNeonGreen was inserted into a predicted flexible loop distal to the membrane-binding surface (Figure 3A and B). This allowed us to explore the location of Rho3 in living cells.

**Figure 3.**
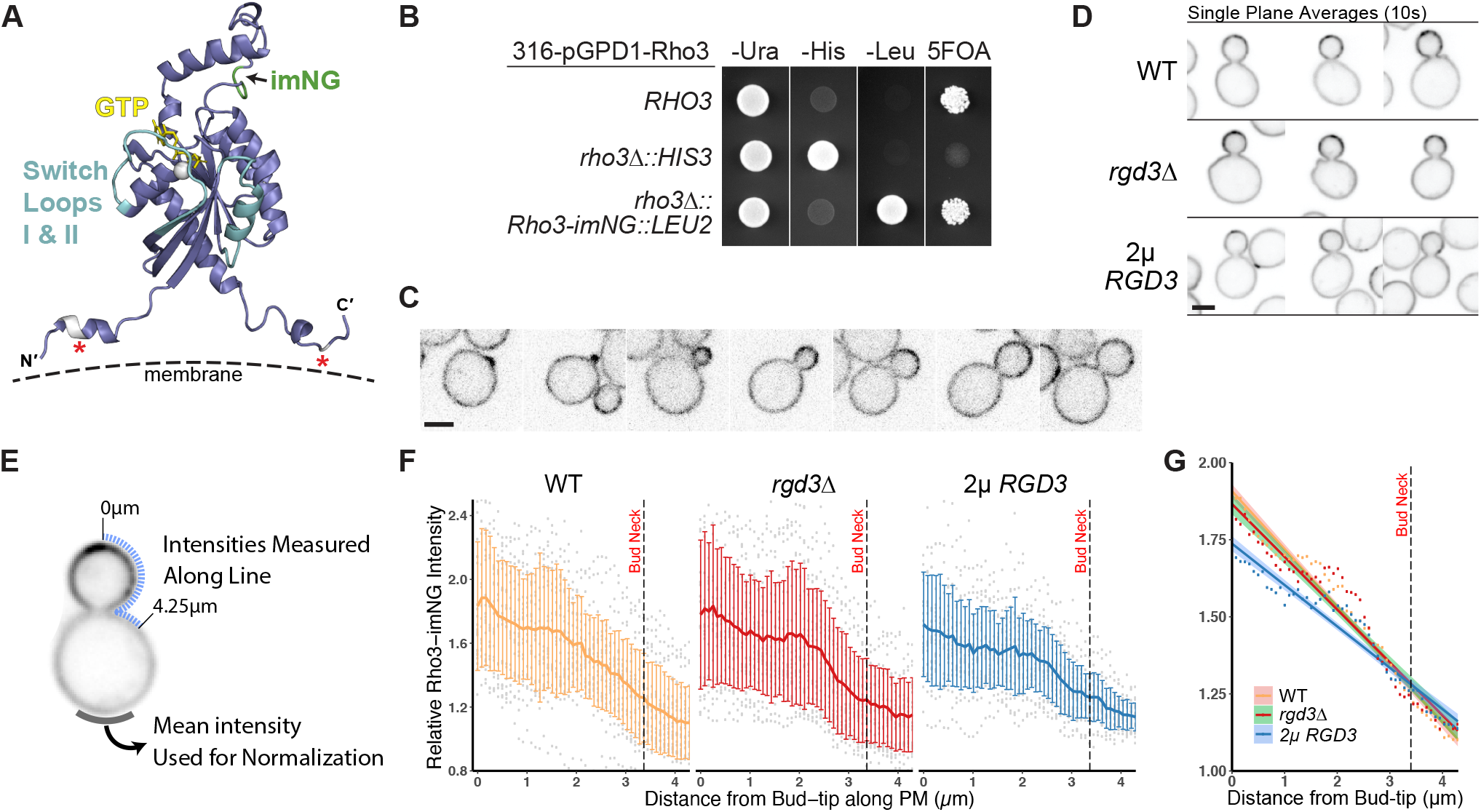
Generation of a functional Rho3 fluorescent reporter and effects of Rgd3 on Rho3 localization. **A.** Modeled 3D structure of Rho3. Modification of both termini (asterisks) is required for functional Rho3 and membrane localization. mNeonGreen was inserted into a predicted flexible loop on the membrane-distal side of Rho3 (imNG). Structure modeled with Robetta (Song et al., 2013). **B.** Rho3-internal-mNeonGreen (imNG) is functional as the sole copy of Rho3 integrated at the endogenous genomic locus. Growth of *rho3::HIS3* or *rho3::Rho3-imNG::LEU2* strains harboring a *URA3-RHO3* plasmid. Eviction of the *URA3* plasmid by selection on 5-FOA shows that Rho3-imNG is functional. **C.** Rho3-imNG localization in various stages of bud-growth; single plane optical sections. Bar, 2μm. **D.** Examples of Rho3-imNG distribution in response to either deletion or overexpression of *RGD3*. Images displayed are single plane, cross-sectional averages of 10s videos. Bar, 2μm. **E.** Schematic representation of the method for measuring distribution and polarity of Rho3-imNG. **F.** Raw data (grey) and average distribution of Rho3-imNG intensity (±SD) along the plasma membrane for each genotype. Intensities measured along dashed lines as in E. *n*=23 per genotype, equal sized buds chosen. **G.** Overlaid linear regression models of data in F ±95%CI. [Video 1 of Rho3-imNG through growth. Maximum projection video timelapse over 3 hours. Images taken every 5 minutes.] [Video 2 of Rho3-imNG overexpression shows internal vesicles. Single plane video. 2μ overexpression is variable and surrounding cells in this video happened to be lower intensity.]

Rho3 internal-mNeonGreen (Rho3-imNG) is predominantly localized to the plasma membrane and enriched at the bud-tip (Figure 3C). Enrichment at the bud-tip remains through mid-sized buds and slowly becomes more evenly distributed in larger buds. Rho3 signal then briefly polarizes to the bud neck just before cytokinesis after which Rho3 localization fluctuates and repolarizes to the membrane proximal to the bud-scar, restarting the cycle (Video 1). During growth, short-lived patches of increased Rho3 concentration can be seen on the plasma membrane, but we did not explore the nature of this phenomenon. While Rho3-imNG could not be visualized on single vesicles when expressed from its endogenous promoter within the genome, overexpression of Rho3-imNG from the strong *GPD1* promoter showed many vesicles, consistent with earlier IF studies (Video 2; Robinson et al., 1999).

To quantitate the overall distribution of Rho3 on the plasma membrane, we performed line scan measurements of the fluorescence intensity along the membrane in medium-sized buds (bud diameter 2.07±0.13 μm) from the bud tip to just past the bud neck over a distance of 4.25μm. The curves were normalized to the mean intensity at the distal region of the mother cells (Figure 3E). This analysis revealed a relatively smooth gradient from the peak intensity at the bud-tip decreasing towards the baseline intensity of the mother. This distribution was not significantly affected in *rgd3*Δ cells but was shallower in cells overexpressing *RGD3* (Figure 3D, F, and G). These findings indicate that Rgd3 can influence the distribution of Rho3 and supports the contention that it functions as a GAP for Rho3.

### Rgd3 localizes to polarized puncta distinct from secretory vesicles

Given Rho3’s prominent localization to the plasma membrane, we next wanted to see if Rgd3 appeared to colocalize with it. Rgd3 tagged C-terminally with mNeonGreen (Rgd3-mNG) is functional, as its overexpression can suppress the conditional *myo2-14 smy1-15* mutant (Figure 4A), thereby allowing us to determine the localization of Rgd3-mNG expressed from its chromosomal locus. *In vivo* imaging revealed dynamic and polarized Rgd3-mNG puncta that were highly concentrated in small budded cells, more distinct in larger budded cells, and often closely associated with the plasma membrane (Figure 4B, Video 3). Imaging the puncta is challenging, as we estimate, based on the fluorescent standard Cse4-mNG, that each punctum contains about 7 molecules of Rgd3-mNG (Figure 4C). Because of limitations imposed by imaging, we cannot determine the lifetime of the Rgd3 puncta, but we suspect they are very short-lived, probably just a few seconds.

**Figure 4.**
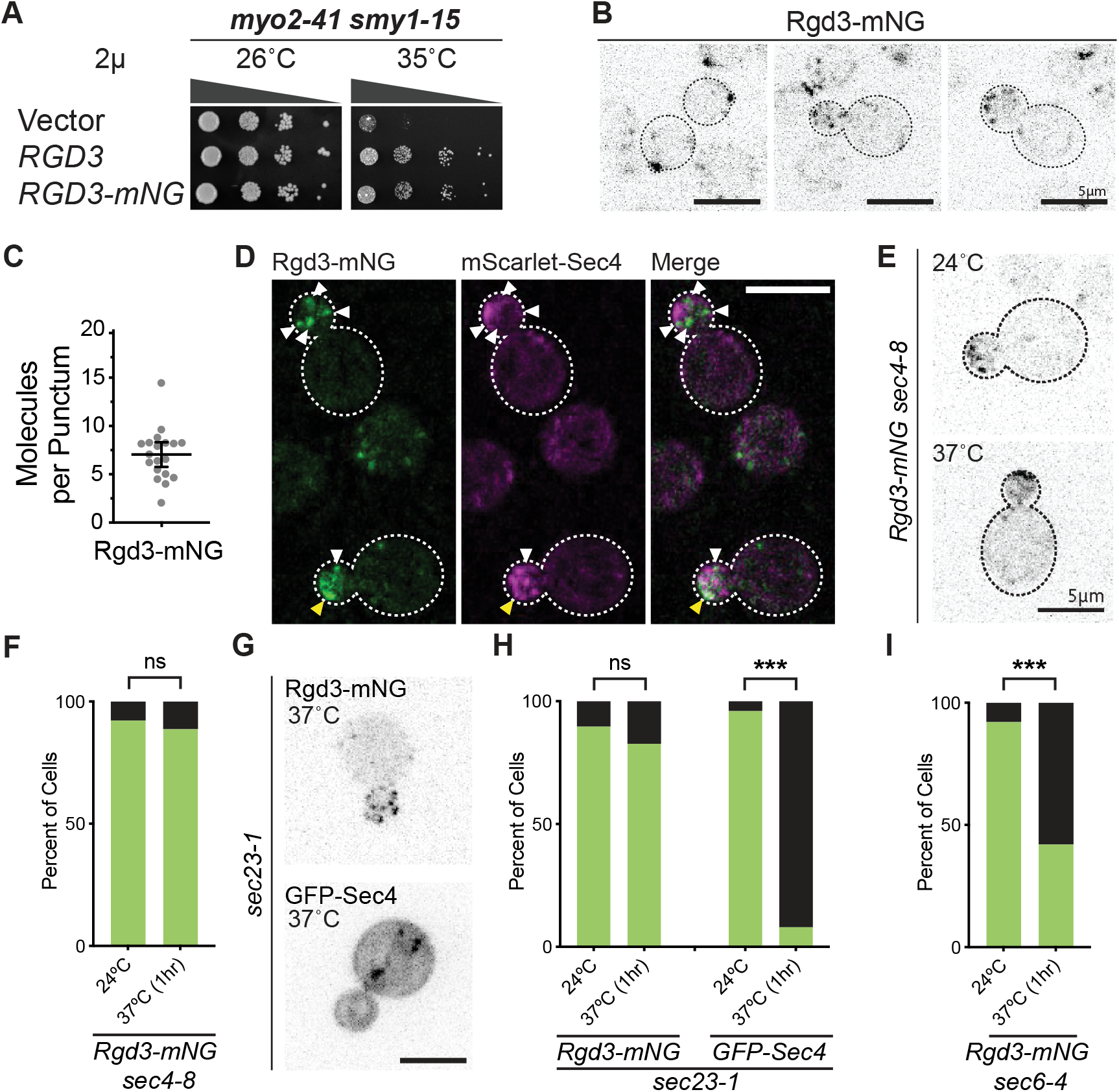
Rgd3 localizes to polarized cytosolic puncta independent of a functional secretory pathway. **A.** Rgd3-mNG is functional in suppression of *myo2-41 smy1-15* temperatures sensitivity **B.** Endogenously tagged Rgd3-mNG localizes to polarized puncta in. **C.** An average of 7±3 Rgd3-mNG molecules are present per punctum (*n*=19). **D.** Single puncta of Rgd3-mNG (white arrows) and mScarlet-Sec4 do not co-localize, although coincidental overlap may occur near the membrane (yellow arrow). **E.** Rgd3-mNG puncta remain polarized in cells where secretory vesicles are depolarized (*sec4-8* cells at 37°C for 1 hour). **F.** Quantification of the presence of polarized Rgd3-mNG in *sec4-8* cells. **G.** Rgd3-mNG puncta remain polarized, while GFP-Sec4 vesicles are abrogated, in cells where the early secretory pathway is blocked (*sec23-1* cells at 37°C for 1 hour). **H.** Quantification of the presence of polarized Rgd3-mNG and GFP-Sec4 puncta/vesicles in *sec23-1* cells. **I.** Quantification of Rgd3-mNG localization in *sec6-4* cells. All quantifications include *n*≥100 cells per condition. ***, p≤0.001 by Z-test for 2 population proportions. Bars, 5μm. [Video 3 of Rgd3-mNG] [Video 4 of fixed Rgd3-mNG mScarlet-Sec4]

The polarized puncta initially appear very similar to diffusely moving secretory vesicles, so we co-imaged Rgd3-mNG and mScarlet-Sec4 and found that the two proteins rarely appeared to colocalize on single puncta (Figure 4D). This exclusivity was further confirmed with fixed cells (Video 4). Thus, the Rgd3 puncta are either a small subset of traditional secretory vesicles or are entirely distinct from them. To test if these puncta are dependent on secretory vesicles, we examined the location of Rgd3-mNG in two conditional mutants that result in either secretory vesicle depolarization (*sec4-8*; Salminen and Novick, 1987) or loss of secretory vesicles by imposing an early secretory block from the endoplasmic reticulum (*sec23-1*; Kaiser and Schekman, 1990). After shifting *sec4-8* cells to their restrictive temperature for one hour, Rgd3-mNG was still present in polarized puncta, indicating that polarized secretory vesicles are not necessary for the polarization of Rgd3 puncta (Figure 4E and F). In *sec23-1* cells, upon shifting to the restrictive temperature, Rgd3-mNG again remained in polarized puncta, whereas GFP-Sec4, as a secretory vesicle marker, was no longer visible on discrete vesicles (Figure 4G and H). We also examined the localization of Rgd3-mNG in *sec6-4* cells which have a conditional defect in secretory vesicle-plasma membrane tethering. In these cells, upon shifting to the restrictive temperature, secretory vesicles are known to accumulate heavily in the bud. Somewhat confoundingly, Rgd3-mNG puncta became moderately less polarized or disappeared altogether (Figure 4I). Collectively, the data reveal that Rgd3 is not located on constitutive secretory vesicles and its polarized distribution is not directly dependent on the secretory pathway.

### Rgd3 associates directly with membranes

The punctate nature of Rgd3’s distribution suggests it may be associated with membranes. Lysates of cells expressing Rgd3-mNG were fractionated on self-forming iodixanol density gradients, a technique developed to separate distinct membrane compartments and previously used to show RabGAP association with membranes (Figure 5A) (Du and Novick, 2001). In these gradients, while Rgd3 floated to a low-density position that indicates vesicle association, this position was repeatedly at a higher density than the majority of secretory vesicles marked by Sec4 (Figure 5B and C). Thus, Rgd3 appears to be membrane associated and on a vesicular compartment largely, or completely, distinct from secretory vesicles.

**Figure 5.**
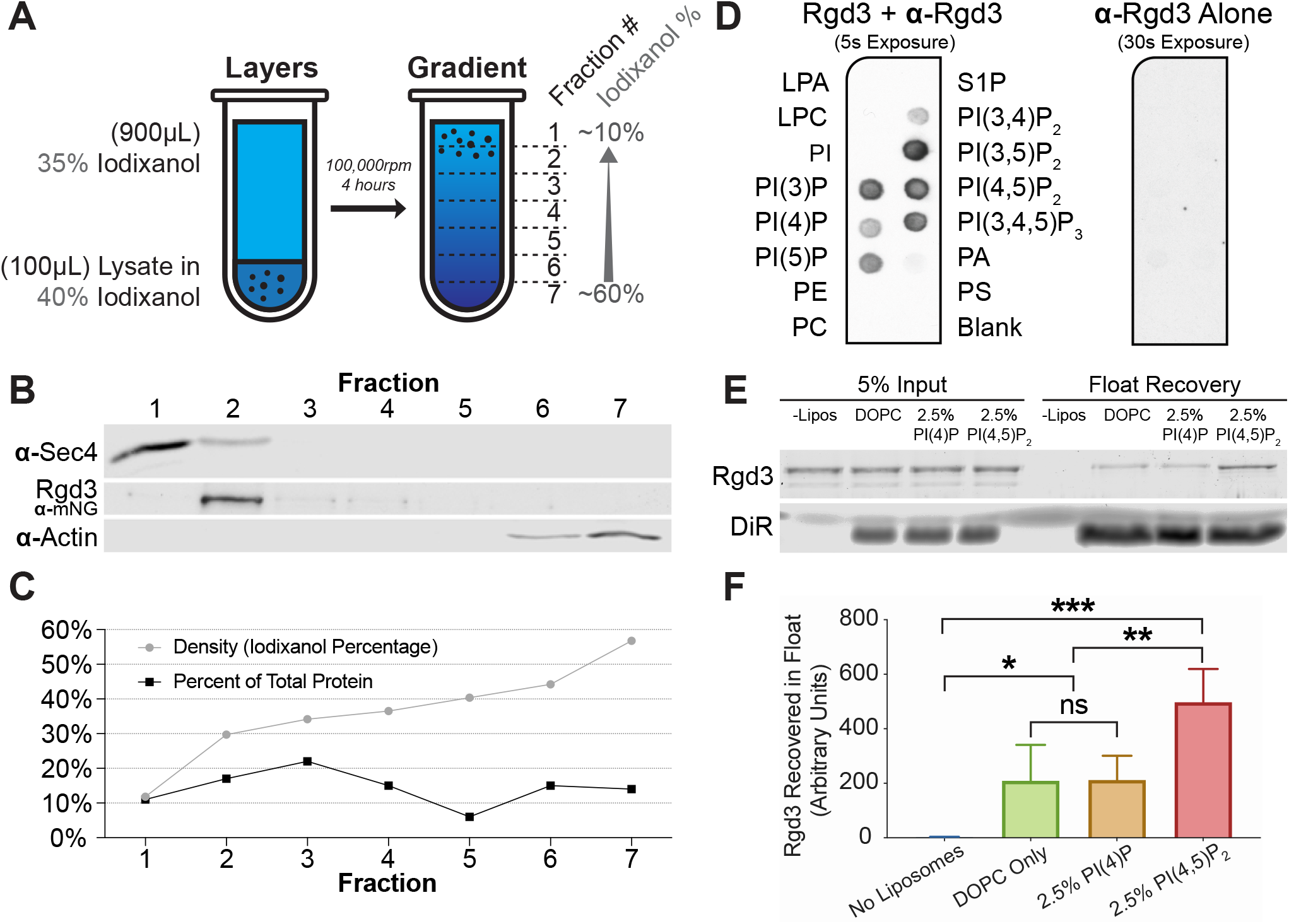
Rgd3 associates with vesicles and binds membranes through PIPs. **A.** Schematic representation of the iodixanol membrane floatation experiment. **B.** Rgd3-mNG floats to a denser fraction than the majority of Sec4 vesicles in a self-forming iodixanol gradient. Representative of 4 gradients. **C.** Iodixanol percentage (grey) and percent of total protein (black) for each fraction from the above experiment. **D.** PIP-strip incubated with 1μg/ml Rgd3 and blotted with rabbit anti-Rgd3. Control strip incubated only with rabbit anti-Rgd3. **E.** Coomassie of liposome-bound Rgd3 from a representative liposome flotation assay. Prepared liposomes contain either DOPC alone, DOPC with 2.5% mol PI(4)P, or DOPC with 2.5% mol PI(4,5)P_2_. Liposomes were labeled with 1% mol DiR dye. **F.** Quantification of Rgd3 recovery in liposome flotation assays. Results from 5 experiments displayed as mean ± SD.

Rgd3 has a predicted F-BAR domain, a structure that generally facilitates direct association with membranes through recognition of phosphatidylinositol-containing lipids (reviewed in Ahmed et al., 2010). As an initial test, full length Rgd3 was expressed in and purified from bacteria (Figure S5), and its ability to associate with specific lipids was tested by blotting on ‘PIP strips’. This revealed a preferential association of Rgd3 with PI(4,5)P_2_ or PI(3,5)P_2_ and varying degrees of affinity for other phosphatidylinositol-phosphate species (Figure 5D). This lipid association was not simply due to negative charge, as no binding to phosphatidylserine was observed. Given its polarized distribution near the plasma membrane, two of the most likely physiologically relevant lipids are PI(4)P and PI(4,5)P_2_. We therefore examined whether purified Rgd3 would associate with artificial liposomes and if its association is influenced by the presence of these regulatory lipids. In floatation assays, Rgd3 associated weakly with DOPC and 2.5% PI(4)P liposomes, and this association was greatly enhanced by the replacement of PI(4)P with 2.5% PI(4,5)P_2_ (Figure 5 E and F). These results mirror the affinities of PI(4)P and PI(4,5)P_2_ seen on the PIP strip and we conclude that Rgd3 has the ability to bind directly to membranes.

### Rgd3 vesicle polarity is dependent on the actin cytoskeleton and Myo2

We next explored whether Rgd3 polarization is dependent on the actin cytoskeleton. Rgd3 is highly polarized in cells induced to shmoo by α-factor treatment, a phenomenon first observed in high-throughput studies (Kraus et al., 2017; Lu et al., 2018). Treatment of shmooing Rgd3-mNG cells with latrunculin B, to depolymerize actin filaments, resulted in complete depolarization of Rgd3-mNG within 40 seconds (Figure 6A). Depolarization of Rgd3 puncta via latrunculin B treatment can also be observed in non-shmooing, nascent and early budding cells (unpublished). Rgd3 polarity, therefore, depends on a dynamic aspect of the actin cytoskeleton.

**Figure 6.**
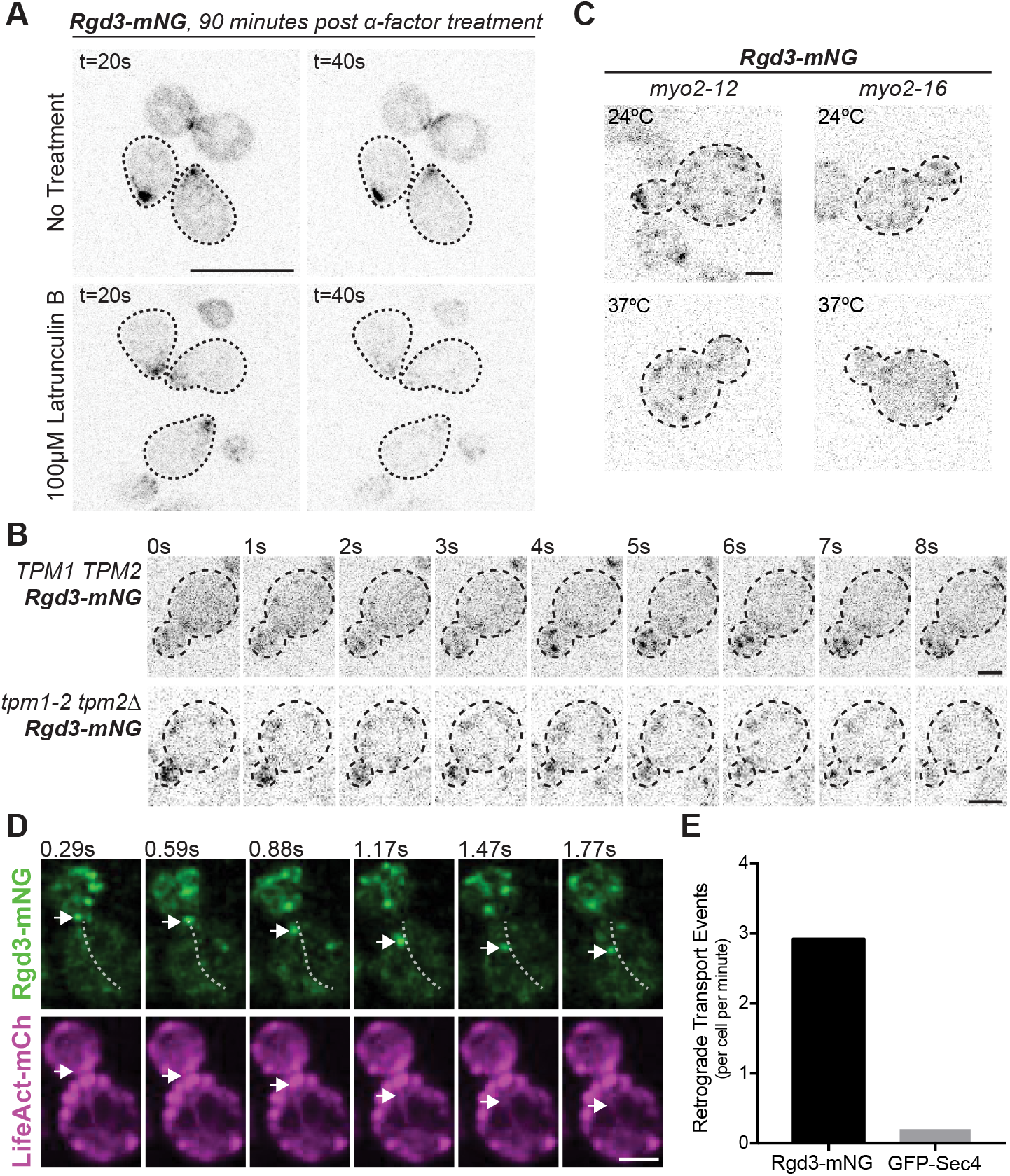
Rgd3 vesicle localization and directed movement is actomyosin dependent. **A.** Rgd3 vesicles, which are highly polarized in alpha-factor-induced shmooing cells, are rapidly depolarized following actin filament depolymerization (100mM Latrunculin B). Images have been equally adjusted for the intense signal at the shmoo-tip; diffuse single vesicles are washed-out. Bar, 10μm. **B.** Rgd3 vesicles depolarize following the destabilization of actin cables. *Rgd3-mNG* or *Rgd3-mNG tpm1-2 tpm2*Δ cells were incubated at room temp and rapidly shifted to a restrictive temperature of 37°C while on the microscope. Imaging began (t=0s) approximately 45s after shift to allow adjustment of optics. **C.** *myo2*^*ts*^ alleles interfere with Rgd3 polarization at both permissive and restrictive temperatures. Bar, 2μm. **D.** Rgd3 vesicles are capable of retrograde (white arrow) movement along actin cables. Actin cable seen in LifeAct-mCherry is traced with a dashed line in the Rgd3-mNG channel. Bar, 2μm. **E.** Quantification of retrograde transport frequency for Rgd3 and Sec4 vesicles. Retrograde transport was defined as 5 consecutive frames of linear motion away from the bud-tip. *n*≥50 cells per genotype. [Video 5 of Rgd3-mNG LifeAct-mCherry]

Latrunculin disrupts all actin filament-dependent processes, including endocytosis. To determine whether Rgd3 polarity is dependent on actin cables, we utilized the temperature sensitive *tpm1-2 tpm2*Δ strain which rapidly destabilizes actin cables, leaving actin cortical patches unscathed (Pruyne et al., 2004a). Indeed, Rgd3-mNG signal was lost from the bud within ~60s of shift to the restrictive temperature (Figure 6B).

As Rgd3 was identified in a strain defective in Myo2-dependent secretory vesicle polarization, a functional link between Rgd3 and Myo2 seems likely. To assess this possibility, we observed Rgd3-mNG localization in two *myo2*^*ts*^ strains: *myo2-12* and *myo2-16*, both of which show a conditional defect in secretory vesicle polarization (Schott et al., 1998). Interestingly, in both of these strains Rgd3 polarity appeared diminished, with more puncta appearing diffuse in the mother cell, even at the permissive temperature (Figure 6C). When raised to the restrictive temperature for an hour, Rgd3 polarization to the bud tip was abolished in *myo2-12* and punctate localization of Rgd3-mNG appeared completely lost in *myo2-16.* These data suggest that Rgd3 vesicle polarity, and maybe function, is dependent on Myo2.

Live-cell imaging of Rgd3-mNG in cells also expressing the F-actin marker LifeAct-mCherry showed that Rgd3 puncta are highly dynamic and occasionally move in a retrograde direction (Figure 6D, Video 5), away from the bud tip and along actin cables, a phenomenon previously observed for the formin Bni1 and occasionally endocytic vesicles (Buttery et al., 2007; Kaksonen et al., 2005). This contrasts with secretory vesicles marked by GFP-Sec4 which almost never move in retrograde (Figure 6E). Since the Rgd3-marked vesicles appear to be unrelated to secretory vesicles, we explored whether they might be downstream of another actin-dependent process, endocytosis.

### Rgd3 is not stably associated with endocytic vesicles

Extensive studies have shown that endocytic vesicles are internalized by Arp2/3-dependent assembly of F-actin at cortical sites (Weinberg and Drubin, 2012), a process that can be imaged using Abp1-mCherry (Shin et al., 2018; Kaksonen et al., 2003). Co-imaging of Abp1-mCherry and chromosomally tagged Rgd3-mNG revealed largely independent localizations and that Rgd3 vesicles are far more dynamic than endocytic patches (Figure S4A, Video 6). Since both Abp1-mCherry and Rgd3-mNG are polarized to the bud, some cases of colocalization do occur. Whether these colocalizations are fortuitous or rather represent a short association of Rgd3-mNG with a subset of endocytic events could not be distinguished. Fixed cells again showed effectively no overlap in Rgd3-mNG and Abp1-mCherry signal (unpublished). Cells compromised for endocytosis show enhanced sensitivity to the toxic arginine analog canavanine as the arginine permease becomes enriched on the cell surface (Lin et al., 2008). However, no consistent phenotype was observed on canavanine for deletion or overexpression of *RGD3* (Figure S4B). Polarized Rgd3 vesicles still appeared in *end3*Δ cells where endocytosis is compromised (Figure S4C) though the frequency and distribution often appeared different from wild-type (unquantified, Video 7). Thus, Rgd3 does not associate with forming endocytic vesicles, but Rgd3 vesicles appear affected by a defect in endocytosis, suggesting some type of relationship.

### Rgd3 contributes to overall polarity and its function is regulated by phosphorylation

Absent a clear pathway responsible for the generation of Rgd3 vesicles, we decided to revisit the original *myo2 smy1* mutant. Tagging of the endogenous Rho3 with imNG in *myo2-41 smy1-15* had little to no effect on growth at room temperature and permitted RGD3 overexpression suppression at 35°C, like the untagged strain (Figure 7A). Fluorescent examination of Rho3-imNG in *myo2-41 smy1-15* cells either overexpressing *RGD3* or containing an empty vector illustrated that Rho3 distribution is largely wild-type at 26°C (Figure 7B). Upon shift to 35°C, however, *myo2-41 smy1-15 Rho3-imNG* accumulates cells with unusual Rho3 polarity and gross morphological defects including, but not limited to: buds growing on buds, buds growing in unusual shapes, and nascent buds forming at bud necks prior to cytokinesis (Fig 7B). Under the same conditions, but when *RGD3* was overexpressed, Rho3-imNG appeared normally polarized and fewer than 3% of cells exhibited similar morphological defects (Figure 7C). These results are consistent with a role for Rgd3 in guiding normal Rho3 distribution on the plasma membrane and bud tip, thereby promoting overall cell polarity and growth.

**Figure 7.**
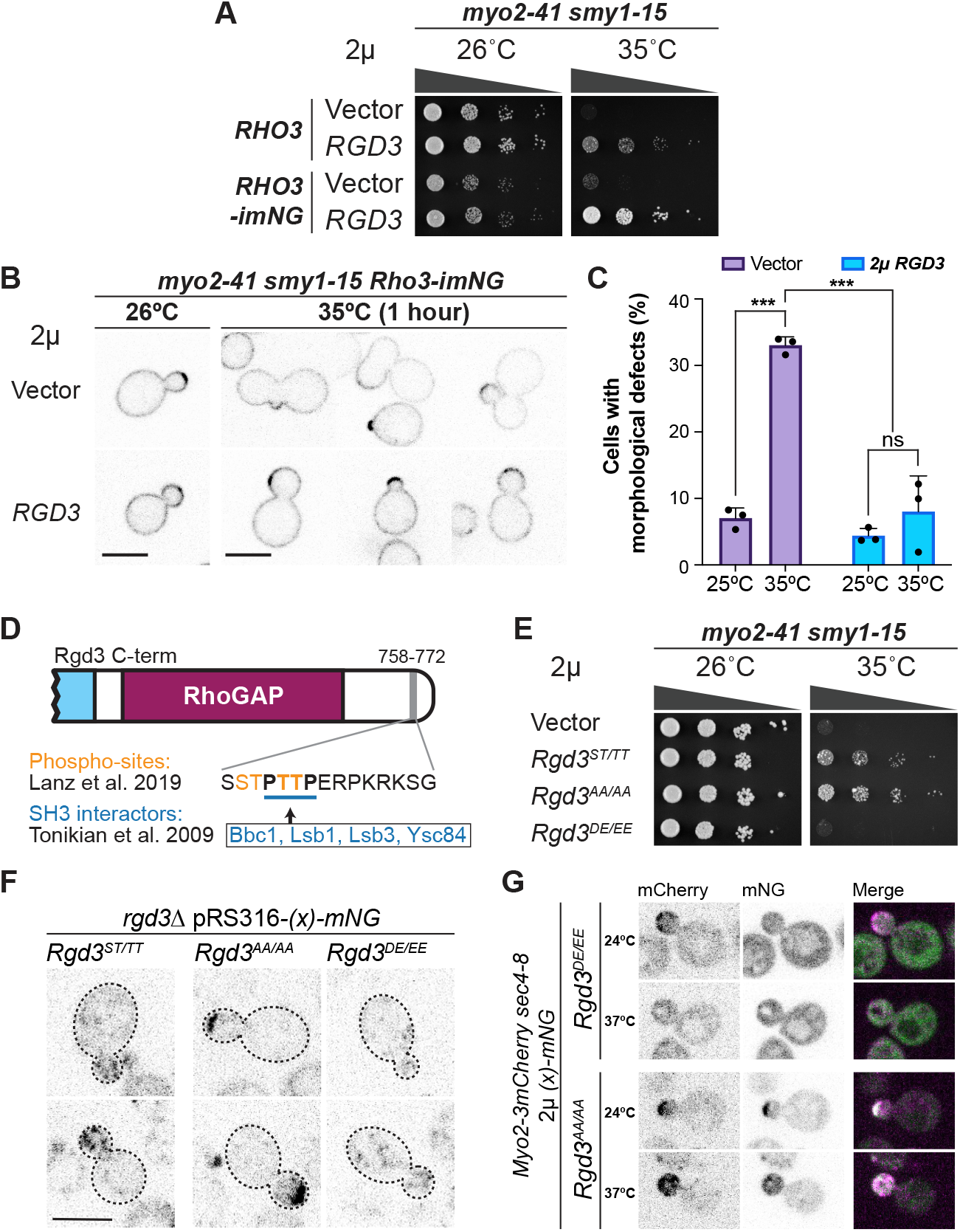
Rgd3 is regulated by C-terminal phosphorylation and participates in endocytosis. **A.** *RGD3* overexpression is capable of suppressing *myo2-41 smy1-15 Rho3-imNG* temperature sensitivity. **B.** *myo2-41 smy1-15 Rho3-imNG* cells exhibit gross morphological defects and mislocalized Rho3 at 35°C which is corrected by overexpression of *RGD3.* **C.** Quantification of cells from B. Data from 3 independent cultures, *n*≥150 cells per culture per condition. ***, p≤0.001 by two-way ANOVA with Sidak’s multiple comparisons test. **D.** The tail of Rgd3, contains a motif (bold/underlined) reported to bind SH3 domains which also contains several phosphorylation sites (orange). **E.** The phospho-dead Rgd3 (*Rgd3*^*AA/AA*^), but not the phospho-mimetic Rgd3 (*Rgd3*^*DE/EE*^), is capable of rescuing *myo2-41 smy1-15*. **F.** *Rgd3*^*AA/AA*^-*mNG* and *Rgd3*^*DE/EE*^-*mNG* mutants localize differently.

Earlier high-throughput work has shown that Rgd3 contains an SH3-domain binding motif at its C-terminus capable of being recognized by the SH3-domain containing factors Bbc1, Lsb1, Lsb3, and Ysc84, all of which participate in actin organization in endocytosis (Figure 7D) (Tonikian et al., 2009). Moreover, the serine and threonines flanking the proline residues of this motif have been found to be phosphorylated in a recent yeast whole phospho-proteome analysis (Lanz et al., 2019). We therefore explored the phenotype of cells in which the serine and threonine residues in Rgd3 were mutated to either alanines (AA/AA mutant) to preclude phosphorylation or aspartic and glutamic acids (DE/EE mutant) to mimic phosphorylation.

Overexpression of the AA/AA mutant was able to suppress the *myo2-41 smy1-15* temperature sensitivity even better than the wild-type ST/TT allele, whereas the DE/EE mutant could not suppress at all (Figure 7E). Further, in *rgd3*Δ cells expressing the Rgd3^DE/EE^-mNG mutant, puncta were very poorly polarized and Rgd3 puncta in cells expressing the Rgd3^AA/AA^-mNG mutant were much more highly polarized (Figure 7F). Since Rgd3-tail phospho-state appears to regulate Rgd3 vesicle polarity and Rgd3 vesicle polarization is dependent on actin cables and Myo2 function, we sought to determine whether Rgd3 was capable of activating and polarizing Myo2 from a depolarized state independent of other polarity cues.

In actively budding cells, Sec4-bound secretory vesicles are the most abundant cargo transported by Myo2 (Pruyne et al., 2004b). For this reason, temperature-sensitive *sec4-8* strains, when raised to the restrictive temperature, results in the inability of Sec4 to activate Myo2 for vesicle transport and a corresponding depolarization of both Myo2 and secretory vesicles (Donovan and Bretscher, 2012). If Rgd3 activates Myo2 to promote Rgd3 vesicle polarization, then introduction of excess *Rgd3*^*AA/AA*^ to *sec4-8 Myo2-3mCherry* cells should permit Myo2 polarization at the restrictive temperature.

In fact, this is exactly what was found. Critically, overexpression of *Rgd3*^*DE/EE*^, the allele that depolarized Rgd3 vesicles, did not exhibit the same polarity-promoting effect on Myo2 as *Rgd3*^*AA/AA*^ (Figure 7G). We conclude that Rgd3 function and localization is negatively regulated by C-terminal phosphorylation.

## Discussion

Yeast has six Rho proteins, three of which, Cdc42, Rho1, and Rho3, are essential, or nearly essential. Cdc42 is the master regulator of polarity (Chiou et al., 2017), Rho1 regulates the PKC pathway as well as the cell wall integrity and maintenance pathways (Levin, 2005), and Rho3 is involved in actin regulation, the establishment of cell polarity, and exocytosis (Robinson et al., 1999; Adamo et al., 1999; Imai et al., 1996; Matsui and Toh-E, 1992). Rho proteins are negatively regulated by RhoGAPs, and the yeast genome encodes 11 putative RhoGAP proteins, 10 of which have now been described, including Rgd3. There must therefore be significant redundancy among RhoGAP proteins, making it unsurprising that none of them individually perform an essential function.

Three of the RhoGAP proteins, Rgd1, Rgd2, and Rgd3 also contain an F-BAR domain that implies they likely associate with moderately curved membranes. Rgd1, a RhoGAP for Rho3 and Rho4, has been shown to localize to polarized vesicles consistent with post-Golgi secretory vesicles (Lefèbvre et al., 2012; Doignon et al., 1999). Rgd2, a RhoGAP for Cdc42 and Rho5, localized to the bud tip in a high-throughput genomic screen (Dubreuil et al., 2018; Roumanie et al., 2001). Here we show that Rgd3 is a RhoGAP for Rho3 and Cdc42, and that it is associated with polarized vesicles. We also found that Rgd3 associated with membranes directly in a phosphoinositide-dependent manner.

While a specific function for Rho3 has not yet been defined, it has been implicated in the establishment of cell polarity, and genetic interactions imply a supporting role in exocytosis through interactions with the exocyst complex. Further, Rho3 has been reported to interact with the lever arm of Myo2 (Forsmark et al., 2011; Robinson et al., 1999). The potential importance of this particular interaction is, at present, unknown, since Rho3 has curiously not been detected (by immunofluorescence or mass spectrometry) on secretory vesicles in cells where it is not overexpressed or mutated (Forsmark et al., 2011; Robinson et al., 1999).

As Rgd3 is a RhoGAP for Rho3, we set out to investigate Rho3’s *in vivo* localization. Both N-terminal and C-terminal tagged Rho3 are non-functional, so we explored internal sites in the protein and found that placement of a fluorescent protein in a predicted flexible loop of chromosomal Rho3 did not affect growth of the cells. A deeper literature search later found that a similar method of internally tagging Cdc42 was employed in *S. Pombe* (Bendezú et al., 2015). Our Rho3-imNG construct allowed us to establish the *in vivo* localization of Rho3, which is normally on the plasma membrane in a gradient decreasing from the bud tip towards the mother cell. Further, this gradient is altered in cells overexpressing Rgd3, supporting the notion that it is a regulator of Rho3.

It is believed that Rho3 is always associated with membranes as it has a bidentate membrane association through both termini and it is unable to be extracted by GDI (Tiedje et al., 2008). Perhaps the most astonishing finding with regards to Rho3 localization is that it is not readily observed on secretory vesicles, despite its established role in enhancing secretion. Further, vesicles observed in cells overexpressing Rho3-imNG did not appear to be particularly polarized, as would be expected from Sec4-marked secretory vesicles. The presence of Rho3 on a different, minor class of vesicles may help explain why it was not identified as a constituent of secretory vesicles.

We next endeavored to define the cellular function and localization of Rgd3. Endogenously tagged Rgd3 clearly localizes to short-lived vesicles which are primarily polarized to the growing bud in an actin cable and Myo2-dependent manner, however, the identity and origin of these vesicles is still unclear. Conditional blocks in secretory vesicle transport (*sec4-8*) and the early secretory pathway (*sec23-1*) had no immediate effect on Rgd3 vesicle abundance or polarity. Reported interactions with the SH3 domain proteins Lsb1, Lsb3, Ysc84, and Bbc1 suggest a role downstream of endocytosis. However, Rgd3 localization appeared minimally changed despite the complete block of endocytosis in *end3*Δ cells (Raths et al., 1993; Tuo et al., 2013). That neither blocks of secretion nor endocytosis drastically alter Rgd3 localization suggests that it may reside on vesicles in a step common to both pathways (e.g. transient recycling endosomes) or some other inter-organellar transport vesicle.

Although we could not establish a clear, single pathway that Rgd3 participates in, a return to the conditional mutant in which it was identified yielded results that support a role of Rgd3 in regulating Rho3 polarity during growth. Overexpression of *RGD3* in *myo2-41 smy1-15 Rho3-imNG* cells eliminates the apparent defect in Rho3 localization and ensuing overall morphology defects seen in an empty vector control. We further showed that the ability of Rgd3 to suppress *myo2-41 smy1-15,* as well Rgd3 vesicle polarity, is dependent on the phosphorylation state of its SH3-interaction motif.

Collectively our data indicate (a) that Rgd3 vesicles are polarized in a Myo2 and actin cable-dependent manner, and also undergo retrograde transport, presumably treadmilling along actin cables; (b) Rgd3 vesicles are not a short-lived intermediate of Sec4-marked secretory vesicles, or a short-lived intermediate of bulk endocytic vesicles; (c) Rgd3 has an F-BAR domain and binds liposomes in a PIP2-enhanced manner, consistent with originating from the plasma membrane; (d) Rgd3 vesicles contribute to Rho3 distribution in the plasma membrane and morphogenesis in a sensitized background, (e) Rgd3 is negatively regulated by C-terminal phosphorylation, and (f) it contains poly-proline motifs that have been shown to bind factors involved in endocytosis.

These properties suggest a speculative model (Figure 8) in which a minor population of regulatory vesicles modulates Rho3 distribution. While we were unable to detect this directly, the simplest explanation for this regulation is that a small population of Rho3 resides on these temporary vesicles, possibly carried internally through endocytosis, where they undergo brief retrograde movement followed by recycling to sites of growth by Myo2. Mutations or backgrounds that result in a large accumulation of Rgd3 vesicles may be needed to test this model fully. It is likely that Rgd3 phosphorylation contributes to its association with the cytoskeletal polarization machinery, so that during highly polarized growth Rgd3 is hypophosphorylated and becomes more phosphorylated as the cell switches to more isotropic growth. As loss of Rgd3 or its over-expression in otherwise wildtype cells has no overt phenotype, testing this model will be challenging and require analysis in genetic backgrounds where its loss causes a strong phenotype.

**Figure 8.**
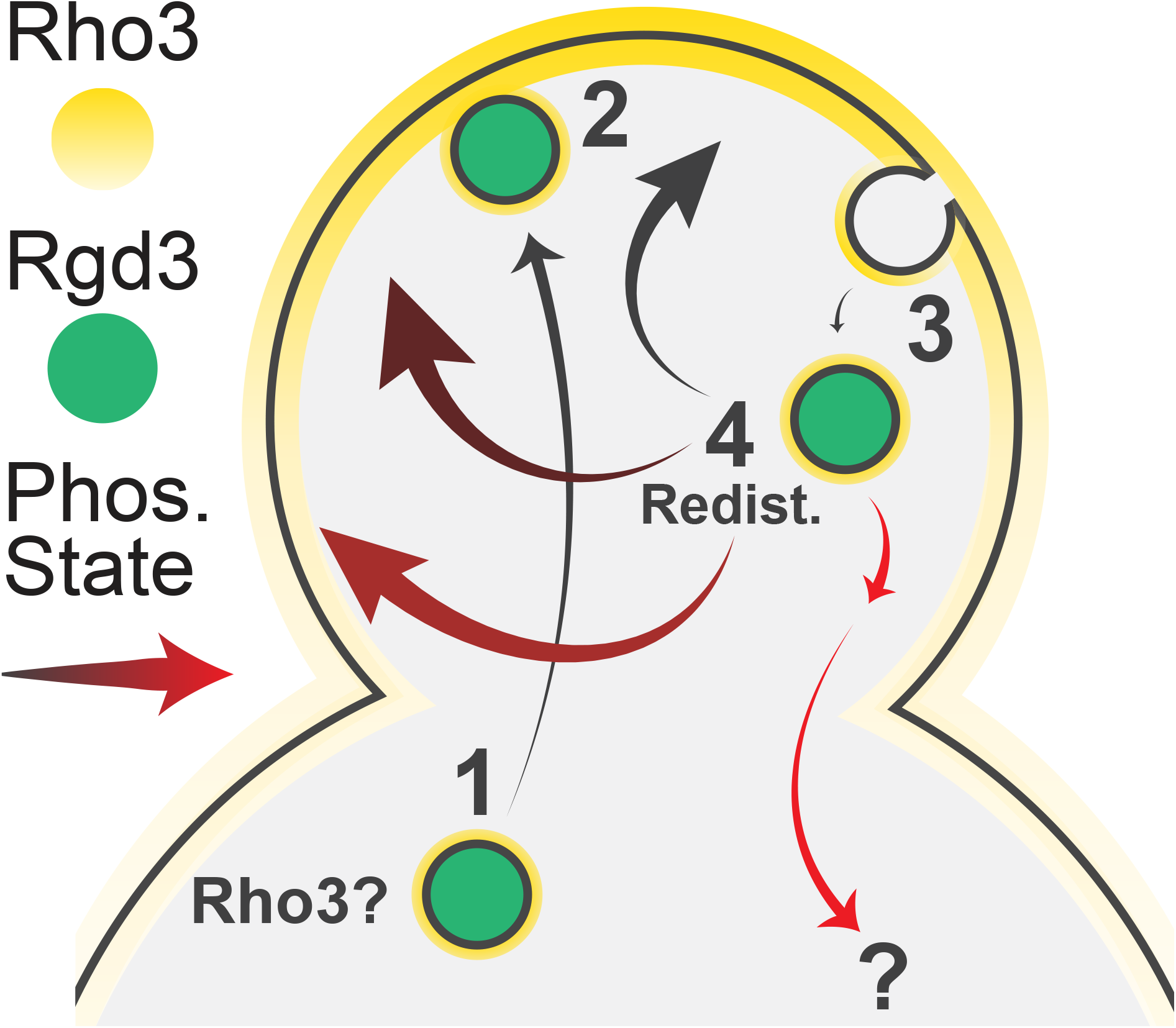
Speculative model of Rgd3 function. **A.** (1) During polarized growth, hypophosphorylated Rgd3 enables activation of Myo2 and the resulting polarization of Rgd3 vesicles, which likely carrying small quantities of Rho3 from the mother cell. (2) Once at sites of growth, Rgd3 vesicles modulate Rho3 localization, potentially by delivery of Rho3 to the membrane via exocyst dependent fusion. (3) Following inevitable endocytosis of Rho3, whether incidental or stimulated, Rgd3 associates with the vesicles to inactivate the internal pool of Rho3. (4) Phosphorylation state of Rgd3 determines how Rho3 is redistributed, with greater phosphorylation likely being associated with more isotropic growth and a less polarized Rho3 distribution, as seen in large buds. Retrograde vesicles may permit cargo degradation or be a spontaneous occurrence.

Our model accounts for why neither complete blockage of endocytosis nor acute blockage of exocytosis abolishes Rgd3 vesicle localization. Furthermore, since unphosphorylated Rgd3 is polarized, overexpression of wild-type Rgd3 would likely result in an increased abundance of unphosphorylated, polarized Rgd3 vesicles. The ability of over-expressed unphosphorylated Rgd3 to polarize Myo2 in the absence of functional secretory vesicles which are its main activator, reveals that Rgd3 can directly, or indirectly, activate Myo2. This ability appears to be responsible for suppression of the defects in secretory vesicle transport in the *myo2 smy1* conditionally sensitive strains. As our data suggest Rgd3 vesicles are distinct from secretory vesicles, its effect on secretory vesicle transport as seen in the conditional mutant likely only occurs upon over-expression. Additionally, Rgd3 over-expression might also contribute by indirectly affecting polarity, morphology, or actin dynamics within the bud.

Finally, a potential clue to the cellular function of Rgd3 in the context of a wild-type cell may lie in one of the other interesting suppressors of our original screen, *SMI1*. Smi1 is reported to be involved in cell wall integrity and synthesis regulation as well as various stress responses (Martin-Yken et al., 2003; Basmaji et al., 2005; Yoshikawa et al., 2008). High throughput studies that included deletion of *YHR182W* imply Rgd3 may also be involved in stress responses and microarray studies suggest that *RGD3* expression is regulated downstream of Hog1, Pho85, and filamentous growth (Yoshikawa et al., 2008; O’Rourke and Herskowitz, 2004; Madhani et al., 1999; Carroll et al., 2001). Further studies will be necessary to determine the mechanistic details of the pathway(s) in which Rgd3 resides.

## Acknowedgements

We would like to acknowledge Cornell colleagues Alan Sulpizio with reagents and advice on the malachite green assay, Carolyn M Highland for assistance on the liposome floatation experiment, and Emr lab members for general advice and reagents. This work was supported by NIH Grants RO1GM39066 and R35GM131751.

## Materials and Methods

### Yeast strains, growth, and transformation

Yeast strains used in this study are listed in Table S1. Standard media and techniques for yeast growing and transformation were used (Sherman, 1991). Gene deletion and chromosomal GFP tagging were performed by standard PCR-mediated techniques (Longtine et al., 1998). Plasmids from the initial screen were isolated from yeast using a genomic DNA isolation kit (Zymo). Dilution assays were performed by first growing cultures in appropriate media to mid-log phase, diluting all to an OD_600_ of 0.3, and plating 10-fold serial dilutions of this on indicated plates. All dilution assays were performed ≥ 3 times.

### cDNA Library screening for overexpression suppressors of myo2 smy1 mutants

1μg of the GAL1-cDNA library was transformed into an overnight culture (5ml) (OD_600_ = ~ 0.8) (Liu et al., 1992). After incubation at 42°C for 45 minutes, the transformants were washed with dH2O once, transferred to a 15ml conical tube and resuspended in 5ml of dH2O. 100μl aliquots were spread on SGal-Ura plates and incubated at 26°C overnight to allow for cDNA expression, and then incubated for 5 days at 35°C. In addition, 100μl of the transformants were spread on to one SC-Ura plate and incubated at 26°C for 5 days to serve as a control for the number of transformants. 80,000 transformants were screened against the *myo2-41 smy1-15* mutant, and 150,000 transformants against the *myo2-57 smy1-12* mutant. The plasmids were recovered from the transformants and sequenced to identify the cDNA.

### DNA constructs

Plasmids used in this study are listed in Table S1. The integrating plasmid pRS306-GFP-Sec4 used to tag Sec4 has been previously described (Donovan and Bretscher, 2012). For constructs utilizing mScarlet, a yeast codon-optimized version of mScarlet (Bindels et al., 2016) was synthesized by IDT for downstream PCR amplification and cloning. Centromeric Rho3-imNG plasmids were generated via Gibson Assembly and genomic integration was done by amplification of Rho3-imNG-LEU2 from pRS415-Rho3-imNG plasmids using primers containing homology to the endogenous Rho3 promotor and 3′UTR. Integration and replacement of endogenous Rho3 was verified by PCR. Site-directed mutagenesis of plasmids was performed via inverse PCR using non-overlapping 5′-phosphorylated primers, one of which contained the intended altered bases at the 5′-end. All plasmid and oligonucleotide sequences are available upon request.

### Microscopy techniques

All micrographs in the main text were acquired on a CSU-X spinning disc confocal microscopy system (Yokogawa, Intelligent Imaging Innovations) with a DMI6000B microscope (Leica), 100×1.45 NA objective (Leica), and an Evolve 512Delta EMCCD (Photometrics) with a 2x magnifying lens (Yokogawa) for a final resolution of 0.084μm/pixel. All images and videos shown are transverse, single plane cross-sections, unless stated otherwise. For live-cell imaging of yeast, cells in mid-log phase were adhered to a glass-bottomed dish (CellVis) coated with ConcanavalinA (EY laboratories) and washed with respective cell medium. Molecule counting for Rgd3 was performed by comparison to Cse4 puncta intensity at anaphase, as described previously (Donovan and Bretscher, 2012). Imaging at elevated temperatures was performed in an environmental chamber (Okolab) following 1-hour incubations in a 37°C water bath, except for the *tpm1-2 tpm2*Δ experiment which was performed in a CherryTemp chamber (Cherry Biotech) for rapid temperature shift. Images were analyzed and processed with Slidebook 6.0 software (Intelligent Imaging Innovations) or FIJI. Images and figures were assembled in Illustrator (Adobe).

### Yeast two-hybrid constructs and analysis

Full length Rgd3 (1-785aa) was fused with the GAL4 activation domain in pGADT7 vector between XmaI and XhoI. Each of all Rho proteins (Rho1, Rho2, Rho3, Rho4, Rho5, Cdc42) were fused with GAL4 DNA binding domain in pGBKT7 vector between XmaI and PstI without their C-terminal CXC box, to prevent prenylation. The AH109 strain co-transformed with both plasmids was selected in media lacking leucine and tryptophan (SC-2DO: double dropout). Interaction was detected by growth on medium lacking leucine, tryptophan and histidine (SC-3DO: triple dropout) or SC-3DO + 1mM 3AT.

### Purification of GST- and 6His-SUMO-tagged proteins

Full length Rho proteins (Rho1, Rho3, and Cdc42) were tagged with GST, while Rgd3, the Rgd3 GAP domain (residues 495-725), and mNeonGreen were tagged with 6His-SUMO. Constructs were expressed in Rosetta 2(DE3) pLysS cells. Cells were grown in terrific broth with antibiotics at 37°C until OD_600_=~1.3. A final concentration of 1mM IPTG was added to induce protein expression overnight at 28°C (Rho proteins), for 5 hours at 28°C (Rgd3/Rgd3-GAP), or overnight at 18°C (mNeonGreen). GST-Rho proteins were isolated on glutathione beads and eluted with excess glutathione. Bacterially expressed 6His-SUMO fusions were isolated on Ni-NTA resin (Qiagen) and eluted from the resin by direct cleavage of SUMO by Ulp1. Proteins were dialyzed into appropriate assay buffers as necessary.

### Antibody production

Rgd3 and mNeonGreen were gel purified following initial SUMO purification and sent to Pocono Rabbit Farm and Laboratory, PA for antibody production. Rabbit antisera against mNeonGreen was affinity purified using mNeonGreen coupled to CnBr beads (MilliporeSigma).

### Malachite green GTPase activity assay

GST-Rho proteins were first dialyzed into Rho Protein Buffer (50mM HEPES, pH7.5, 100mM KCl, 1mM DTT) to remove excess phosphate and diluted to equal concentration following a Bradford protein assay. Rgd3-GAP was dialyzed into malachite reaction buffer (50mM HEPES, pH7.5, 100mM KCl, 5mM EDTA, 10mM MgCl_2_). To prepare malachite green reagent, 30ml of 0.045% (w/v) malachite green oxalate (Alfa Aezea #A16186) was combined with 20ml of 4.2% (w/v) ammonium molybdate in 4M HCL and nutated for 30 minutes before filtering through a 0.2μm filter. Tween20 was added to 0.02% before use to prevent precipitation.

The malachite green assay for measuring phosphate has been described previously (Maehama et al., 2000). In a 96-well plate held on ice, each Rho protein was suspended to a final concentration of 25μM in nucleotide exchange buffer (50mM Tris, pH7.5, 250mM KCl, 5mM EDTA, 10mM MgCl_2_) with or without 100μM freshly prepared GTP. Reactions were incubated at 30°C for 30 minutes before MgCl_2_ was spiked-in to a final concentration of 10mM to halt nucleotide exchange. Rgd3-GAP was then added to a final concentration equimolar to that of Rho (for stimulated GTP-hydrolysis) or a similar volume of malachite reaction buffer was added (for intrinsic hydrolysis). Reactions were again incubated at 30°C for 30 minutes. For readout of released phosphate, 100μL of malachite green reagent was added and allowed to react for 5 minutes at 37°C. Reactions were equilibrated to room temperature for 5 minutes before being quickly read for absorbance at 620nm on a 96-well plate reader. Phosphate released was calculated relative to the absorbance of a simultaneously prepared and read standard curve of sodium phosphate. Each reaction was performed in triplicate, with appropriate triplicate controls for each reagent and protein alone.

### Measurement of Rho3-imNG distribution

Mid-sized cells (~2μm diameter buds) expressing Rho3-imNG at the endogenous locus were captured in approximately 10 second videos, focused on the bud neck. The resulting frames of each video were averaged in FIJI to yield representative images of Rho3 distributions while minimizing the contributions of temporary punctate structures on the plasma membrane. Freehand lines with a width of 4 pixels were manually drawn from the apparent bud tip (determined by perpendicularity to the bud neck, not signal intensity) through approximately 0.5μm of the mother membrane. The point of inflection at the bud neck was noted along the intensity profile plot and used for later alignment of the data. Intensity profiles were normalized to the average intensity of a ~0.5μm stretch of the antipolar membrane in the mother to account for variations in Rho3 expression. Data was analyzed in Open Source R Studio.

### Iodixanol gradients and floatation assay

Iodixanol floatation assays were performed essentially as described in Du and Novick (2001), except that gradients were centrifuged in a TLA-100.2 rotor at 100,000rpm for 4 hours. Protein content per fraction was measured via Bradford assay and iodixanol density was measured via absorbance at 244nm (Schröder et al., 1997).

### Immunoblotting

Samples from iodixanol floatation experiments were resolved by SDS-PAGE, transferred to Immobilon-P membrane (MilliporeSigma), then blocked for 1 hour in 5% milk in TBS-T (0.1% Tween). Membranes were first incubated with primary affinity purified rabbit anti-mNG and visualized with secondary HRP-goat anti-rabbit. Blots were probed again with primary mouse anti-actin (MilliporeSigma) and rabbit anti-Sec4. After washing, membranes were incubated with secondary AlexaFluor-conjugated goat anti-mouse and donkey anti-rabbit antibodies. Membranes were finally analyzed using the Odyssey infrared imaging system (Li-COR).

### PIP-Strip

Membranes spotted with 100pmol of various lipids (PIP-Strip, Echelon Biosciences) were blocked overnight at 4°C in blocking buffer (PBS, 0.1% Tween20, 3% BSA). Membranes were incubated at room temperature for 1 hour with purified 1μg/ml full-length Rgd3 in blocking buffer, washed 3x with PBS-T, and then incubated with rabbit anti-Rgd3 antisera for another hour. Negative controls were performed using PIP-strips that had not been incubated with purified Rgd3 but were also blotted with rabbit anti-Rgd3 antisera. Strips were again washed, incubated with secondary HRP-goat anti-rabbit for 1 hour and visualized on X-ray film with Amersham ECL detection reagents (GE).

### Liposome Extrusion

Liposomes were generated as previously described (Richardson and Fromme, 2015). Briefly, individual lipid stocks (Avanti Polar Lipids, Inc.) were combined in 96.5:2.5:1 DOPC:PIP:DiR-dye molar ratios in the presence of chloroform and methanol. Lipid films were vacuum-dried in pear-shaped flasks and rehydrated overnight to 1mM at 37°C in HK buffer (20mM HEPES pH 7.4 and 150mM KOAc). Lipids were extruded in a mini-extruder (Avanti Polar Lipids, Inc) with 100-nm filters (Whatman) using 25 passes through the filter. Liposomes were stored at 4°C and were generally used within 1 week of extrusion.

### Discontinuous Sucrose Density Gradient

Liposome floatation assays were performed essentially as previously described (Richardson and Fromme, 2015). For liposome-binding, 6μg of full-length Rgd3 protein was combined with 20μl of 1mM liposomes and brought to a final volume of 80μl in HK buffer. This mixture was allowed to incubate for 1 hour at room temperature before gently, but thoroughly mixing in 50μl of 2.5M sucrose in HK buffer. Unbound protein was separated from liposomes by transferring 100μl of this mixture to 7×20mm PC ultracentrifuge tubes (Beckman) and carefully layering an equal volume of 0.75M sucrose buffer followed by 25μl of sucrose-free HK buffer, and gradients were spun 100,000rpm for 20min at 20°C in a TLA-100 ultracentrifuge rotor (Beckman). Top layers were collected, and bound protein was assessed by SDS-PAGE stained with QC Colloidal Coomassie (BioRad). Lipid recovery was normalized to the input samples by measuring relative DiR fluorescence on an Odyssey Imager (Li-COR).

### Tetrad dissection

Genetic interaction between *rgd3*Δ and *rgd1*Δ was tested by dissecting tetrads after sporulation. Cells with opposite mating types were grown together overnight at 26°C for mating. Mating efficiency was checked by shmoo projection and zygote formations the next day under the light microscope. Cells were streaked onto diploid-selective plates (SC-Met/-Lys) and single resulting colonies were grown in YPD overnight, washed with PBS, and resuspended in sporulation media (1% yeast extract, 1% potassium acetate, and 0.05% glucose). Sporulation was carried out over 5-7 days. Once tetrads had formed, 25μl of zymolyase (stock 1mg/ml) was added to 50μl of sporulated cells. The digestion was incubated at 37°C for 6-7 minutes. Tetrads were dissected using an MSM Singer instrument with a 25μm fiber needle.

---

Figure S1. **Replication of key experiments in *myo2-57 smy1-12* and effect of *RGD3* overexpression or *rgd3*Δ on wild-type cells.**

**A.** Assessment of overexpression suppression of the identified cDNAs on 2μ vectors in *myo2-57 smy1-12* cells.

**B.** *RHO3* and *CDC42* 2μ overexpression exhibit the greatest counter-suppression of temperature sensitivity in *myo2-57 smy1-12* 2μ *RGD3.*

**C.** Neither deletion nor overexpression of *RGD3* exhibit growth defects in wild-type cells.

Figure S2. **Overexpression and localization of *Rgd3*^*R546A*^ compared to wildtype *Rgd3.***

**A.** Western blot with anti-Rgd3 in wildtype, *rgd3*Δ, and cells overexpressing *Rgd3* or *Rgd3*^*R546A*^ via 2μ plasmid. Anti-actin loading control below.

**B.** Localization of Rgd3-mNG and *Rgd3*^*R546A*^-*mNG* expressed in *rgd3*Δ cells via CEN plasmid.

Figure S3. ***rgd3*Δ *rgd1*Δ strains exhibit no apparent synthetic growth or cytoskeletal defects.**

**A.** A *rgd3*Δ *rgd1*Δ strain is viable. Tetrad dissection of *rgd3*Δ*::HisMX* X *rgd1*Δ*::Ura3.* White boxes indicate double delete strains.

**B.** *rgd3*Δ *rgd1*Δ strains have no synthetic growth defects.

**C.** Phalloidin staining of F-actin in WT, *rgd3*Δ, *rdg1*Δ, and *rgd1*Δ *rgd3*Δ exhibits no clear actin-morphological defects.

Figure S4. **Rgd3 is not stably associated with endocytic vesicles and has no direct effect on endocytosis.**

**A.** Localization of Rgd3-MnG and Abp1-mCherry.

**B.** Growth of ten-fold serial dilutions of cells lacking (*rgd3*Δ) or over-expressing *RGD3.*

**C.** Localization of Rgd3-mNG is *end3*Δ cells compromised for endocytosis.

[Video 6 of Abp1-mCherry and Rgd3-mNG illustrates minimal meaningful signal overlap.]

[Video 7 *end3*Δ *Rgd3-mNG* exhibits variable Rgd3 vesicle abundance and distribution.]

Figure S5. **Purified proteins used in biochemical assays.**

Table S1. **List of strains (Table 1) and plasmids (Table 2) used in study.**

## References

Adamo, J.E., G. Rossi, and P. Brennwald. 1999. The Rho GTPase Rho3 Has a Direct Role in Exocytosis That Is Distinct from Its Role in Actin Polarity. Mol Biol Cell. 10:4121–4133. doi:10.1091/mbc.10.12.4121.

Basmaji, F., H. Martin-Yken, F. Durand, A. Dagkessamanskaia, C. Pichereaux, M. Rossignol, and J. Francois. 2005. The ‘interactome’ of the Knr4/Smi1, a protein implicated in coordinating cell wall synthesis with bud emergence in Saccharomyces cerevisiae. Mol Genet Genomics. 275:217–230. doi:10.1007/s00438-005-0082-8.

Bendezú, F.O., V. Vincenzetti, D. Vavylonis, R. Wyss, H. Vogel, and S.G. Martin. 2015. Spontaneous Cdc42 Polarization Independent of GDI-Mediated Extraction and Actin-Based Trafficking. Plos Biol. 13:e1002097. doi:10.1371/journal.pbio.1002097.

Beningo, K.A., S.H. Lillie, and S.S. Brown. 2000. The Yeast Kinesin-related Protein Smy1p Exerts Its Effects on the Class V Myosin Myo2p via a Physical Interaction. Mol Biol Cell. 11:691–702. doi:10.1091/mbc.11.2.691.

Bindels, D.S., L. Haarbosch, L. van Weeren, M. Postma, K.E. Wiese, M. Mastop, S. Aumonier, G. Gotthard, A. Royant, M.A. Hink, and T.W.J. Gadella. 2016. mScarlet: a bright monomeric red fluorescent protein for cellular imaging. Nat Methods. 14:53–56. doi:10.1038/nmeth.4074.

Bourne, H.R. 1997. The arginine finger strikes again. Nature. 389:673–674. doi:10.1038/39470.

Brockerhoff, S., R. Stevens, and T. Davis. 1994. The unconventional myosin, Myo2p, is a calmodulin target at sites of cell growth in Saccharomyces cerevisiae. J Cell Biology. 124:315–323. doi:10.1083/jcb.124.3.315.

Burchett, S.A. 2002. Regulators of G Protein Signaling. J Neurochem. 75:1335–1351. doi:10.1046/j.1471-4159.2000.0751335.x.

Buttery, S.M., S. Yoshida, and D. Pellman. 2007. Yeast Formins Bni1 and Bnr1 Utilize Different Modes of Cortical Interaction during the Assembly of Actin Cables. Mol Biol Cell. 18:1826–1838. doi:10.1091/mbc.e06-09-0820.

Carroll, A.S., A.C. Bishop, J.L. DeRisi, K.M. Shokat, and E.K. O’Shea. 2001. Chemical inhibition of the Pho85 cyclin-dependent kinase reveals a role in the environmental stress response. Proc National Acad Sci. 98:12578–12583. doi:10.1073/pnas.211195798.

Chiou, J., M.K. Balasubramanian, and D.J. Lew. 2017. Cell Polarity in Yeast. Annu Rev Cell Dev Bi. 33:77–101. doi:10.1146/annurev-cellbio-100616-060856.

Doignon, F., C. Weinachter, O. Roumanie, and M. Crouzet. 1999. The yeast Rgd1p is a GTPase activating protein of the Rho3 and Rho4 proteins. Febs Lett. 459:458–462. doi:10.1016/s0014-5793(99)01293-4.

Du, L.-L., and P. Novick. 2001. Yeast Rab GTPase-activating Protein Gyp1p Localizes to the Golgi Apparatus and Is a Negative Regulator of Ypt1p. Mol Biol Cell. 12:1215–1226. doi:10.1091/mbc.12.5.1215.

Dubreuil, B., E. Sass, Y. Nadav, M. Heidenreich, J.M. Georgeson, U. Weill, Y. Duan, M. Meurer, M. Schuldiner, M. Knop, and E.D. Levy. 2018. YeastRGB: comparing the abundance and localization of yeast proteins across cells and libraries. Nucleic Acids Res. 47:D1245–D1249. doi:10.1093/nar/gky941.

Forsmark, A., G. Rossi, I. Wadskog, P. Brennwald, J. Warringer, and L. Adler. 2011. Quantitative proteomics of yeast post-Golgi vesicles reveals a discriminating role for Sro7p in protein secretion. Traffic Cph Den. 12:740–53. doi:10.1111/j.1600-0854.2011.01186.x.

Hammer, J.A., and J.R. Sellers. 2012. Walking to work: roles for class V myosins as cargo transporters. Nat Rev Mol Cell Bio. 13:13–26. doi:10.1038/nrm3248.

Imai, J., A. Toh-e, and Y. Matsui. 1996. Genetic analysis of the Saccharomyces cerevisiae RHO3 gene, encoding a rho-type small GTPase, provides evidence for a role in bud formation. Genetics. 142:359–69.

Johnston, G.C., J.A. Prendergast, and R.A. Singer. 1991. The Saccharomyces cerevisiae MYO2 gene encodes an essential myosin for vectorial transport of vesicles. J Cell Biology. 113:539–551. doi:10.1083/jcb.113.3.539.

Kaksonen, M., Y. Sun, and D.G. Drubin. 2003. A Pathway for Association of Receptors, Adaptors, and Actin during Endocytic Internalization. Cell. 115:475–487. doi:10.1016/s0092-8674(03)00883-3.

Kaksonen, M., C.P. Toret, and D.G. Drubin. 2005. A Modular Design for the Clathrin- and Actin-Mediated Endocytosis Machinery. Cell. 123:305–320. doi:10.1016/j.cell.2005.09.024.

Kraus, O.Z., B.T. Grys, J. Ba, Y. Chong, B.J. Frey, C. Boone, and B.J. Andrews. 2017. Automated analysis of high-content microscopy data with deep learning. Mol Syst Biol. 13:924. doi:10.15252/msb.20177551.

Lanz, M.C., K. Yugandhar, S. Gupta, E. Sanford, V. Faça, S. Vega, A. Joiner, C. Fromme, H. Yu, and M.B. Smolka. 2019. In-depth and 3-Dimensional Exploration of the Budding Yeast Phosphoproteome. Biorxiv. 700070. doi:10.1101/700070.

Lefèbvre, F., V. Prouzet-Mauléon, M. Hugues, M. Crouzet, A. Vieillemard, D. McCusker, D. Thoraval, and F. Doignon. 2012. Secretory Pathway-Dependent Localization of the Saccharomyces cerevisiae Rho GTPase-Activating Protein Rgd1p at Growth Sites. Eukaryot Cell. 11:590–600. doi:10.1128/ec.00042-12.

Levin, D.E. 2005. Cell Wall Integrity Signaling in Saccharomyces cerevisiae. Microbiol Mol Biol R. 69:262–291. doi:10.1128/mmbr.69.2.262-291.2005.

Lillie, S.H., and S.S. Brown. 1992. Suppression of a myosin defect by a kinesin-related gene. Nature. 356:358–361. doi:10.1038/356358a0.

Lillie, S.H., and S.S. Brown. 1994. Immunofluorescence localization of the unconventional myosin, Myo2p, and the putative kinesin-related protein, Smy1p, to the same regions of polarized growth in Saccharomyces cerevisiae. J Cell Biology. 125:825–842. doi:10.1083/jcb.125.4.825.

Lin, CH, MacGurn, JA, Chu, T, Stefan, CJ, and Emr, SD (2008). Arrestin-Related Ubiquitin-Ligase Adaptors Regulate Endocytosis and Protein Turnover at the Cell Surface. Cell 135, 714–725.

Liu, H., J. Krizek, and A. Bretscher. 1992. Construction of a GAL1-regulated yeast cDNA expression library and its application to the identification of genes whose overexpression causes lethality in yeast. Genetics. 132:665–73.

Longtine, M.S., A.M. III, D.J. Demarini, N.G. Shah, A. Wach, A. Brachat, P. Philippsen, and J.R. Pringle. 1998. Additional modules for versatile and economical PCR-based gene deletion and modification in Saccharomyces cerevisiae. Yeast. 14:953–961. doi:10.1002/(sici)1097-0061(199807)14:10<953::aid-yea293>3.0.co;2-u.

Lu, A.X., Y.T. Chong, I.S. Hsu, B. Strome, L.-F. Handfield, O. Kraus, B.J. Andrews, and A.M. Moses. 2018. Integrating images from multiple microscopy screens reveals diverse patterns of protein subcellular localization change. Elife. 7:e31872. doi:10.7554/elife.31872.

Lwin, K.M., D. Li, and A. Bretscher. 2016. Kinesin-related Smy1 enhances the Rab-dependent association of myosin-V with secretory cargo. Mol Biol Cell. 27:2450–62. doi:10.1091/mbc.e16-03-0185.

Madeira, F., Y.M. Park, J. Lee, N. Buso, T. Gur, N. Madhusoodanan, P. Basutkar, A.R.N. Tivey, S.C. Potter, R.D. Finn, and R. Lopez. 2019. The EMBL-EBI search and sequence analysis tools APIs in 2019. Nucleic Acids Res. 47:W636–W641. doi:10.1093/nar/gkz268.

Madhani, H.D., T. Galitski, E.S. Lander, and G.R. Fink. 1999. Effectors of a developmental mitogen-activated protein kinase cascade revealed by expression signatures of signaling mutants. Proc National Acad Sci. 96:12530–12535. doi:10.1073/pnas.96.22.12530.

Maehama, T., G.S. Taylor, J.T. Slama, and J.E. Dixon. 2000. A Sensitive Assay for Phosphoinositide Phosphatases. Anal Biochem. 279:248–250. doi:10.1006/abio.2000.4497.

Martin-Yken, H., A. Dagkessamanskaia, F. Basmaji, A. Lagorce, and J. Francois. 2003. The interaction of Slt2 MAP kinase with Knr4 is necessary for signalling through the cell wall integrity pathway in Saccharomyces cerevisiae. Mol Microbiol. 49:23–35. doi:10.1046/j.1365-2958.2003.03541.x.

Matsui, Y., and A. Toh-E. 1992. Yeast RHO3 and RHO4 ras superfamily genes are necessary for bud growth, and their defect is suppressed by a high dose of bud formation genes CDC42 and BEM1. Mol Cell Biol. 12:5690–5699. doi:10.1128/mcb.12.12.5690.

O’Rourke, S.M., and I. Herskowitz. 2004. Unique and Redundant Roles for HOG MAPK Pathway Components as Revealed by Whole-Genome Expression Analysis. Mol Biol Cell. 15:532–542. doi:10.1091/mbc.e03-07-0521.

Pruyne, D., L. Gao, E. Bi, and A. Bretscher. 2004a. Stable and Dynamic Axes of Polarity Use Distinct Formin Isoforms in Budding Yeast. Mol Biol Cell. 15:4971–4989. doi:10.1091/mbc.e04-04-0296.

Pruyne, D., L. Gao, E. Bi, and A. Bretscher. 2004b. MECHANISMS OF POLARIZED GROWTH AND ORGANELLE SEGREGATION IN YEAST. Annu Rev Cell Biol. 20(1), 559–591. https://dx.doi.org/10.1146/annurev.cellbio.20.010403.103108

Raths, S., J. Rohrer, F. Crausaz, and H. Riezman. 1993. end3 and end4: two mutants defective in receptor-mediated and fluid-phase endocytosis in Saccharomyces cerevisiae. J Cell Biology. 120:55–65. doi:10.1083/jcb.120.1.55.

Ren, J., L. Wen, X. Gao, C. Jin, Y. Xue, and X. Yao. 2008. CSS-Palm 2.0: an updated software for palmitoylation sites prediction. Protein Eng Des Sel. 21:639–644. doi:10.1093/protein/gzn039.

Richardson, B.C., and J.C. Fromme. 2015. Biochemical methods for studying kinetic regulation of Arf1 activation by Sec7. Methods Cell Biol. 130:101–26. doi:10.1016/bs.mcb.2015.03.020.

Robinson, N.G.G., L. Guo, J. Imai, A. Toh-e, Y. Matsui, and F. Tamanoi. 1999. Rho3 of Saccharomyces cerevisiae, Which Regulates the Actin Cytoskeleton and Exocytosis, Is a GTPase Which Interacts with Myo2 and Exo70. Mol Cell Biol. 19:3580–3587. doi:10.1128/mcb.19.5.3580.

Roumanie, O., C. Weinachter, I. Larrieu, M. Crouzet, and F. Doignon. 2001. Functional characterization of the Bag7, Lrg1 and Rgd2 RhoGAP proteins from Saccharomyces cerevisiae. Febs Lett. 506:149–156. doi:10.1016/s0014-5793(01)02906-4.

Schott, D., J. Ho, D. Pruyne, and A. Bretscher. 1999. The Cooh-Terminal Domain of Myo2p, a Yeast Myosin V, Has a Direct Role in Secretory Vesicle Targeting. J Cell Biology. 147:791–808. doi:10.1083/jcb.147.4.791.

Schott, D., T. Huffaker, and A. Bretscher. 2002. Microfilaments and microtubules: the news from yeast. Curr Opin Microbiol. 5:564–574. doi:10.1016/s1369-5274(02)00369-7.

Schröder, M., R. Schäfer, and P. Friedl. 1997. Spectrophotometric Determination of Iodixanol in Subcellular Fractions of Mammalian Cells. Anal Biochem. 244:174–176. doi:10.1006/abio.1996.9861.

Shin, M., J. van Leeuwen, C. Boone, and A. Bretscher. 2018. Yeast Aim21/Tda2 both regulates free actin by reducing barbed end assembly and forms a complex with Cap1/Cap2 to balance actin assembly between patches and cables. Mol Biol Cell. 29:923–936. doi:10.1091/mbc.e17-10-0592.

Song, Y., F. DiMaio, R.Y.-R. Wang, D. Kim, C. Miles, T. Brunette, J. Thompson, and D. Baker. 2013. High-resolution comparative modeling with RosettaCM. Struct Lond Engl 1993. 21:1735–42. doi:10.1016/j.str.2013.08.005.

Stevens, R.C., and T.N. Davis. 1998. Mlc1p Is a Light Chain for the Unconventional Myosin Myo2p in Saccharomyces cerevisiae. J Cell Biology. 142:711–722. doi:10.1083/jcb.142.3.711.

Tang, K., Y. Li, C. Yu, and Z. Wei. 2019. Structural mechanism for versatile cargo recognition by the yeast class V myosin Myo2. J Biol Chem. 294:5896–5906. doi:10.1074/jbc.ra119.007550.

Tiedje, C., I. Sakwa, U. Just, and T. Höfken. 2008. The Rho GDI Rdi1 regulates Rho GTPases by distinct mechanisms. Mol Biol Cell. 19:2885–96. doi:10.1091/mbc.e07-11-1152.

Tonikian, R., X. Xin, C.P. Toret, D. Gfeller, C. Landgraf, S. Panni, S. Paoluzi, L. Castagnoli, B. Currell, S. Seshagiri, H. Yu, B. Winsor, M. Vidal, M.B. Gerstein, G.D. Bader, R. Volkmer, G. Cesareni, D.G. Drubin, P.M. Kim, S.S. Sidhu, and C. Boone. 2009. Bayesian Modeling of the Yeast SH3 Domain Interactome Predicts Spatiotemporal Dynamics of Endocytosis Proteins. Plos Biol. 7:e1000218. doi:10.1371/journal.pbio.1000218.

Tuo, S., K. Nakashima, and J.R. Pringle. 2013. Role of Endocytosis in Localization and Maintenance of the Spatial Markers for Bud-Site Selection in Yeast. Plos One. 8:e72123. doi:10.1371/journal.pone.0072123.

Weinberg, J., and D.G. Drubin. 2012. Clathrin-mediated endocytosis in budding yeast. Trends Cell Biol. 22:1–13. doi:10.1016/j.tcb.2011.09.001.

Wong, H.C., J. Mao, J.T. Nguyen, S. Srinivas, W. Zhang, B. Liu, L. Li, D. Wu, and J. Zheng. 2000. Structural basis of the recognition of the Dishevelled DEP domain in the Wnt signaling pathway. Nat Struct Biol. 7:1178–1184. doi:10.1038/82047.

Wu, H., and P. Brennwald. 2010. The Function of Two Rho Family GTPases Is Determined by Distinct Patterns of Cell Surface Localization. Mol Cell Biol. 30:5207–5217. doi:10.1128/mcb.00366-10.

Yoshikawa, K., T. Tanaka, C. Furusawa, K. Nagahisa, T. Hirasawa, and H. Shimizu. 2008. Comprehensive phenotypic analysis for identification of genes affecting growth under ethanol stress in Saccharomyces cerevisiae. Fems Yeast Res. 9:32–44. doi:10.1111/j.1567-1364.2008.00456.x.

Zimmermann, L., A. Stephens, S.-Z. Nam, D. Rau, J. Kübler, M. Lozajic, F. Gabler, J. Söding, A.N. Lupas, and V. Alva. 2018. A Completely Reimplemented MPI Bioinformatics Toolkit with a New HHpred Server at its Core. J Mol Biol. 430:2237–2243. doi:10.1016/j.jmb.2017.12.007.

