## Supplementary figures and images for "Yeast Rgd3 is a phospho-regulated F-BAR-containing RhoGAP involved in the regulation of Rho3 distribution and cell morphology"

### Supplemental Figure 1

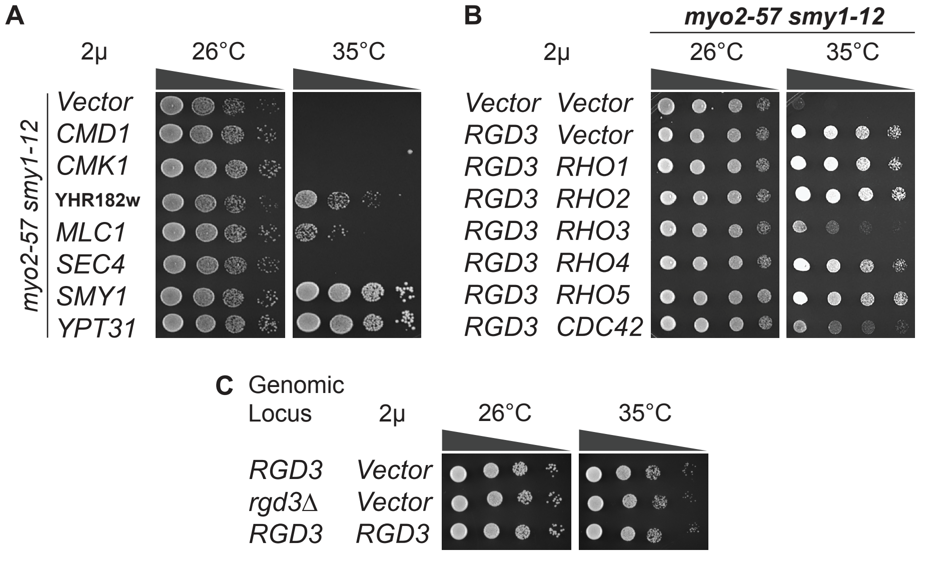

### Supplemental Figure 2

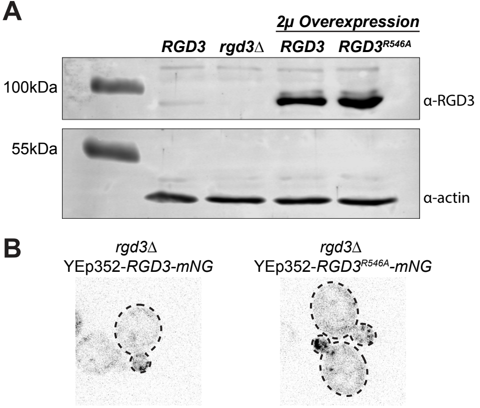

### Supplemental Figure 3

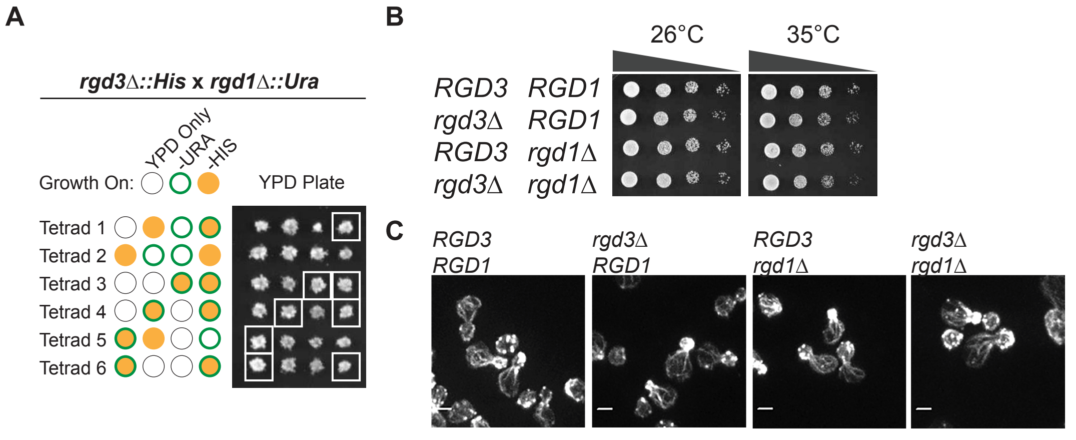

### Supplemental Figure 4

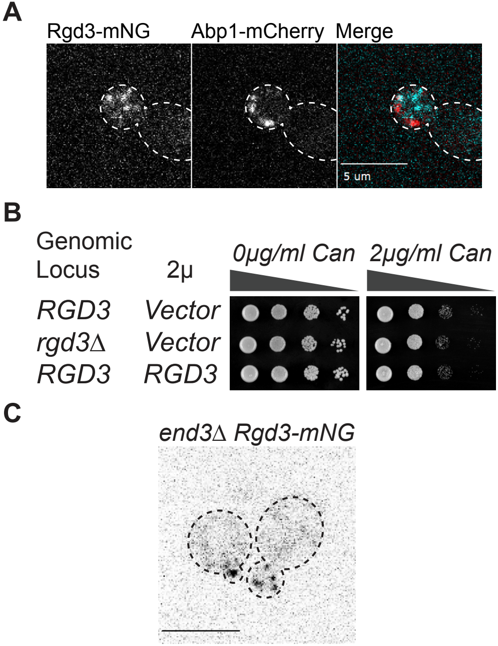
