## Supplemental Tables for "Yeast Rgd3 is a phospho-regulated F-BAR-containing RhoGAP involved in the regulation of Rho3 distribution and cell morphology"

| TABLE 1 | | | |
|---|---|---|---|
| **Strain ID (Alias)** | **Mat Type** | **Genotype** | **Origin** |
| BY4741 | a | *his3Δ1 leu2Δ0 ura3Δ0 met15Δ0* | Brachmann et al., 1998 |
| BY4742 | alpha | *his3Δ1 leu2Δ0 ura3Δ0 lys2Δ0* | Brachmann et al., 1998 |
| ABY3319 | a | *myo2-41::HIS3 trp1Δ::KanMX* | Lwin et al., 2016 |
| ABY4013 | a | *smy1-15::TRP1 trp1Δ::KanMX* | Lwin et al., 2016 |
| ABY3320 | a | *myo2-41::HIS3 smy1-15::TRP1 trp1Δ::KanMX* | Lwin et al., 2016 |
| ABY3316 | a | *myo2-57::HIS3 trp1Δ::KanMX* | Lwin et al., 2016 |
| ABY4012 | a | *smy1-12::TRP1 trp1Δ::KanMX* | Lwin et al., 2016 |
| ABY3351 | a | *myo2-57::HIS3 smy1-12::TRP1 trp1Δ::KanMX* | Lwin et al., 2016 |
| ABY9222 | a | *rgd3Δ(YHR182wΔ)::HIS3* | This Study |
| ABY4711 | alpha | *rgd1Δ::URA3* | This Study |
| ABY4715 (AH109a) | ? | *rgd3Δ::HIS3 rgd1Δ::URA3* | This Study |
| ABY2766 | a | *trp1-901, leu2-3, 112, ura3-52, his3-200, gal4Δ, gal80Δ* | Clontech (TaKaRa Bio) |
| ABY9300 | alpha | *rho3Δ::HIS3 + pRS316-pADH-RHO3* | This Study |
| ABY9315 | alpha | *rho3Δ::RHO3-imNG::LEU2* | This Study |
| ABY9316 | alpha | *rho3Δ::RHO3-imNG::LEU2 rgd3Δ::HIS3* | This Study |
| ABY9127 | alpha | *RGD3-mNG::LEU2* | This Study |
| ABY9200 | a | *RGD3-mNG::URA3* | This Study |
| ABY9201 | alpha | *RGD3-mNG::URA3* | This Study |
| ABY9182 | a | *CSE4-mNG::URA3* | This Study |
| ABY9204 | a | *mScarlet-SEC4::URA3 RGD3-mNG::LEU2* | This Study |
| ABY4045 | alpha | *sec4-8::HIS3* | This Study |
| ABY9298 | alpha | *sec4-8::HIS3 RGD3-mNG::LEU2* | This Study |
| ABY9051 | alpha | *sec6-4::HIS3* | This Study |
| ABY9299 | alpha | *sec6-4::HIS3 RGD3-mNG::LEU2* | This Study |
| ABY9054 | alpha | *sec6-4::HIS3 GFP-SEC4::URA3* | This Study |
| ABY0824 (RSY281) | alpha | *his4-619 ura3-52 sec23-1* | Kaiser and Schekman, 1990 |
| ABY3440 | alpha | *his4-619 ura3-52 sec23-1 GFP-SEC4::URA3* | Donovan and Bretscher, 2012 |
| ABY9337 | alpha | *his4-619 ura3-52 sec23-1 RGD3-mNG::URA3* | This Study |
| ABY9338 | a | *LifeAct-mCherry::HIS3 RGD3-mNG::URA3* | This Study |
| ABY9369 | a | *RGD3-mNG::URA3 ABP1-mCherry::LEU2* | This Study |
| ABY9390 | a | *end3Δ::KanMX RGD3-mNG::URA3* | This Study |
| ABY9391 | a | *myo2-41::HIS3 smy1-15::TRP1 trp1Δ::KanMX RHO3-imNG::LEU2* | This Study |
| ABY6013 | a | *tpm1-2::Leu tpm2Δ::KanMX* | Pruyne, Schott, and Bretscher, 1998 |
| ABY9423 | a | *tpm1-2::Leu tpm2Δ::KanMX Rgd3-mNG::URA3* | This Study |
| ABY9435 | a | *Myo2-mCherry::LEU2 sec4-8::HIS3 YEp352-Rgd3(DE/EE)* | This Study |
| ABY9436 | a | *Myo2-mCherry::LEU2 sec4-8::HIS3 YEp352-Rgd3(AA/AA)* | This Study |
| ABY9437 | a | *Myo2-mCherry::LEU2 sec4-8::HIS3* | This Study |

| **TABLE 2** | | | |
|---|---|---|---|
| **Plasmid ID** | **Vector Backbone** | ***Insert*** | **Origin** |
| **2μ Overexpression** | | | |
| 16 | YEp351 | *empty* | Hill et al., 1986 |
| 14 | YEp352 | *empty* | Hill et al., 1986 |
| 3857 | YEp351 | *SMY1* | Lwin et al., 2016 |
| 3858 | YEp352 | *SMY1* | Lwin et al., 2016 |
| 4097 | YEp351 | *CMD1* | This Study |
| 3952 | YEp352 | *CMD1* | This Study |
| 3953 | YEp351 | *CMK1* | This Study |
| 3863 | YEp351 | ***RGD3 (YHR182w)*** | This Study |
| 3864 | YEp352 | ***RGD3 (YHR182w)*** | This Study |
| 3954 | YEp351 | *MLC1* | This Study |
| 3955 | YEp352 | *MLC1* | This Study |
| 3850 | YEp351 | *SEC4* | This Study |
| 3851 | YEp352 | *SEC4* | This Study |
| 3854 | YEp351 | *YPT31* | This Study |
| 3852 | YEp351 | *YPT32* | This Study |
| 3853 | YEp352 | *YPT32* | This Study |
| 3919 | YEp351 | *RHO1* | This Study |
| 3920 | YEp352 | *RHO1* | This Study |
| 3921 | YEp351 | *RHO2* | This Study |
| 3922 | YEp352 | *RHO2* | This Study |
| 3923 | YEp351 | *RHO3* | This Study |
| 3865 | YEp352 | *RHO3* | This Study |
| 3924 | YEp351 | *RHO4* | This Study |
| 3925 | YEp352 | *RHO4* | This Study |
| 3926 | YEp351 | *RHO5* | This Study |
| 3927 | YEp352 | *RHO5* | This Study |
| 3928 | YEp351 | *CDC42* | This Study |
| 3929 | YEp352 | *CDC42* | This Study |
| 3915 | YEp351 | ***RGD3(R546A)*** | This Study |
| 3916 | YEp352 | ***RGD3(R546A)*** | This Study |
| 3946 | YEp352 | *RGD1* | This Study |
| 4632 | YEp352 | *RGD2* | This Study |
| 4633 | YEp352 | *BEM3* | This Study |
| 4631 | YEp352 | *RGA1* | This Study |
| 3944 | YEp352 | *RGA2* | This Study |
| 3937 | YEp352 | *BAG7* | This Study |
| 3941 | YEp352 | *SAC7* | This Study |
| 3939 | YEp352 | *LRG1* | This Study |
| 4481 | YEp352 | ***RGD3-mNG*** | This Study |
| 4642 | YEp352 | ***RGD3(AA/AA)*** | This Study |
| 4639 | YEp352 | ***RGD3(DE/EE)*** | This Study |
| 3052 | pRS425 | *empty* | Christianson et al., 1992 |
| 4432 | pRS425 | *RHO3* | This Study |
| 4149 | pRS425 | *RHO3(Q74L)* | This Study |
| 4464 | pRS425 | *RHO3(T30N)* | This Study |
| **Centromeric and Integrating** | | | |

| | | | |
|---|---|---|---|
| 3832 | pRS316 | *pADH-RHO3* | This Study |
| 4665 | pRS415 | *pRHO3-RHO3-imNG* | This Study |
| 3964 | pRS303 | *sec4-8* | This Study |
| 3101 | pRS303 | *sec6-4* | This Study |
| 3471 | pRS306 | *GFP-SEC4* | Donovan and Bretscher, 2012 |
| 3833 | pRS305 | *GFP-SEC4* | Lwin et al., 2016 |
| 4103 | pRS306 | *mScarlet-SEC4* | This Study |
| 4196 | pRS303 | *pADH-LifeAct-mCherry* | This Study |
| 4490 | pRS316 | ***RGD3-mNG*** | This Study |
| 4640 | pRS316 | ***RGD3(AA/AA)-mNG*** | This Study |
| 4643 | pRS316 | ***RGD3(DE/EE)-mNG*** | This Study |
| **Yeast 2-Hybrid** | | | |
| 2519 | pGADT7 | *empty* | Clontech (TaKaRa Bio) |
| 2518 | pGBKT7 | *empty* | Clontech (TaKaRa Bio) |
| 3991 | pGADT7 | ***RGD3*** | This Study |
| 3996 | pGBKT7 | *RHO1* | This Study |
| 3997 | pGBKT7 | *RHO2* | This Study |
| 3998 | pGBKT7 | *RHO3* | This Study |
| 3999 | pGBKT7 | *RHO4* | This Study |
| 4000 | pGBKT7 | *RHO5* | This Study |
| 4001 | pGBKT7 | *CDC42* | This Study |
| **Protein Purification** | | | |
| 3836 | pE-SUMO | ***RGD3*** | This Study |
| 3837 | pE-SUMO | ***RGD3 GAP Domain*** | This Study |
| 4409 | pE-SUMO | *mNG* | This Study |
| 3840 | pGEX-6p3 | *RHO1* | This Study |
| 3842 | pGEX-6p3 | *RHO3* | This Study |
| 3845 | pGEX-6p3 | *CDC42* | This Study |
| **Genomic Tagging** | | | |
| 4430 | pFA6a-URA3 | *mScarlet* | This Study |
| All others previously reported | | | |
